# Multi-modal Diffusion Model with Dual-Cross-Attention for Multi-Omics Data Generation and Translation

**DOI:** 10.1101/2025.02.27.640020

**Authors:** Erpai Luo, Lei Wei, Minsheng Hao, Xuegong Zhang, Qiao Liu

## Abstract

Single-cell multi-omics technologies offer unprecedented opportunities to decipher complex cellular mechanisms. To overcome experimental limitations in scale, cost, and coverage, powerful computational methods are essential for integrating diverse data modalities and generating high-fidelity in-silico data. In this paper, we present scDiffusion-X, a latent diffusion model specifically designed for this purpose. The core innovation is a Dual-Cross-Attention (DCA) module that adaptively learns intricate, hidden relationships between different molecular modalities, offering a more flexible and interpretable approach than traditional integration strategies. Extensive benchmarking experiments demonstrate that scDiffusion-X excels at generating realistic multi-omics data, preserving cellular heterogeneity and global data structures with excellent scalability. Distinct from existing multi-omics simulators, scDiffusion-X uniquely enables high-fidelity modality translation by predicting one molecular modality from another and provides robust uncertainty quantification. Beyond data generation, we designed a gradient-based interpretation framework to transform DCA module into a discovery tool, enabling inference of comprehensive cell-type-specific heterogeneous gene regulatory networks (GRNs). By integrating state-of-the-art generative modeling with deep biological interpretability, scDiffusion-X serves as a powerful tool for dissecting regulatory relationships, predicting perturbation responses, poised to accelerate discovery in single-cell multi-omics research.

## 1. Main

Single-cell multi-omics technologies have revolutionized our understanding of cellular diversity by simultaneously profiling various molecular modalities^1,2^, such as the transcriptome, chromatin accessibility, epigenome, and proteome, within individual cells. For instance, technologies such as 10x Multiome and SNARE-seq^3^ simultaneously profile single-cell RNA expression and chromatin accessibility, while approaches like CITE-seq^4^ measure cell surface protein and transcriptome within a single-cell. These advancements provide a more comprehensive and systematic view of cellular states^5^, developmental trajectories^6^, and interactions^7^ than single-modality techniques, opening new opportunities for understanding complex biological systems^8,9^.

Although single-cell multi-omics technologies have made significant progress, challenges exist in acquiring high-quality, large-scale multi-omics datasets due to the need for meticulous sample preparation, and specialized experimental conditions^10,11^. Also, simultaneous measurements of distinct modalities often result in trade-offs between resolution and coverage. Consequently, it is difficult to obtain large quantities of high-quality multi-omics data at scale^12^. Furthermore, multi-omics sequencing remains more expensive than more established single-omics sequencing methods, creating barriers to their widespread application^13,14^.

To mitigate these challenges, computational models have been developed to integrate and generate in-silico multi-omics data. For instance, MultiVI^15^ extends the Variational Autoencoder (VAE)-based scRNA-seq analysis tool scVI^16^ and the scATAC-seq analysis tool peakVI^17^, enabling the joint analysis of multi-omics data and the reconstruction of paired datasets by learning a shared latent space. While effective, VAE-based models often struggle with representing complex, multi-modal distributions and may produce overly smooth data^18-20^. CFGen^21^ employs a flow-based generative model to produce multi-omics data and enhances classifier-free guidance in Flow Matching with multiple attribute instances, thereby improving the conditional generation capabilities of the multi-omics data simulator. However, flow-based models are constrained by invertible transformations, which can limit their expressiveness when modeling highly complex distributions and the ability to elucidate relationships between modalities^22,23^. scDesign3^24^ is a recent statistical model designed to simulate single-cell multi-omics data by learning parameters from observed data. Despite the primary goal of these models is to provide simulation data for benchmarking the computational methods rather than to generate large-scale realistic data for biological analyses, it struggles to scale effectively with large datasets due to their limited capacity to handle the high dimensionality of multi-omics data^25^.

To address the above limitations, we present scDiffusion-X, a deep generative model specifically designed for single-cell multi-omics data. This model integrates a multi-modal latent denoising diffusion probability architecture to generate multi-omics data under user-specific conditions. By leveraging the advanced capabilities of diffusion models, regarded as one of the most advanced deep generative frameworks across various domains^26-29^, scDiffusion-X exhibits exceptional scalability and flexibility. Its iterative denoising process not only ensures the generation of biologically meaningful data but also maintains comprehensive mode coverage, making it uniquely suited for modeling intricate relationships across diverse modalities and handling high-dimensional, large-scale single-cell multi-omics datasets.

A key innovation of scDiffusion-X is the Dual-Cross-Attention (DCA) module, which is designed to enable adaptive and interpretable information exchange between different modalities. Compared to traditional integration strategies such as early integration, late integration, and intermediate integration^30^, which impose fixed interaction patterns (e.g., concatenation), the DCA provides a much more flexible and comprehensive way to integrate multi-modalities information and offers more insights into the intrinsic relationship between different modalities. To enhance interpretability, we proposed a gradient-based interpretability framework with the DCA to identify cell-type specific heterogeneous gene regulatory networks (GRNs), offering a biologically grounded understanding of regulatory interactions.

Unlike the existing simulation methods that only focus on generation by learning the joint distribution of multi-modalities, scDiffusion-X uniquely supports modality translation by simultaneously learning the conditional distribution of one modality based on another. More importantly, our approach incorporates the uncertainty quantification, allowing for probabilistic characterization of generated data. This represents a significant advancement over existing modality translation methods, which typically yield point estimates^31,32^.

To rigorously evaluate the capabilities of scDiffusion-X, we conducted a series of experiments including conditional multi-omics data generation and modality translation. The results demonstrate that scDiffusion-X consistently outperforms the existing methods by generating high-quality multi-omics data across diverse conditions and scales. Furthermore, scDiffusion-X introduces a novel, gradient-based interpretability approach that deciphers the relationships between genes and chromatin peaks to construct cell-type specific heterogeneous GRNs. This unique capability transforms scDiffusion-X from a data simulator into a biologically meaningful framework for understanding gene regulation mechanisms. Overall, these findings highlight the utility of scDiffusion-X as a powerful tool for generative modeling of multi-omics data, enabling the dissection of complex regulatory relationships, prediction of perturbation responses, and identification of potential biomarkers and regulatory patterns.

## 2. Results

### 2.1 Overview of scDiffusion-X model

scDiffusion-X is a multi-modal latent denoising diffusion probability model for single-cell multi-omics data generation. The architecture has two main primary components: a multimodal autoencoder and a multimodal denoising network. The multimodal autoencoder encodes each modality from the multi-omics data into a low-dimensional latent space, facilitating diffusion-based learning, and subsequently decodes it back to its original space (Figure 1a). The multimodal denoising network operates in latent space, progressively refining noisy representations to recover biologically meaningful multi-omics data (Figure 1b-c). A key innovation of scDiffusion-X is the Dual-Cross-Attention (DCA) module, integrated within the multimodal denoising network, to effectively integrate information from different modalities in an interpretable way (Figure 1d). Additionally, condition labels, such as cell type, tissue, disease state, or experimental condition, are embedded into the time-step representations, allowing scDiffusion-X to generate context-specific multi-omics data.

**Figure 1.**
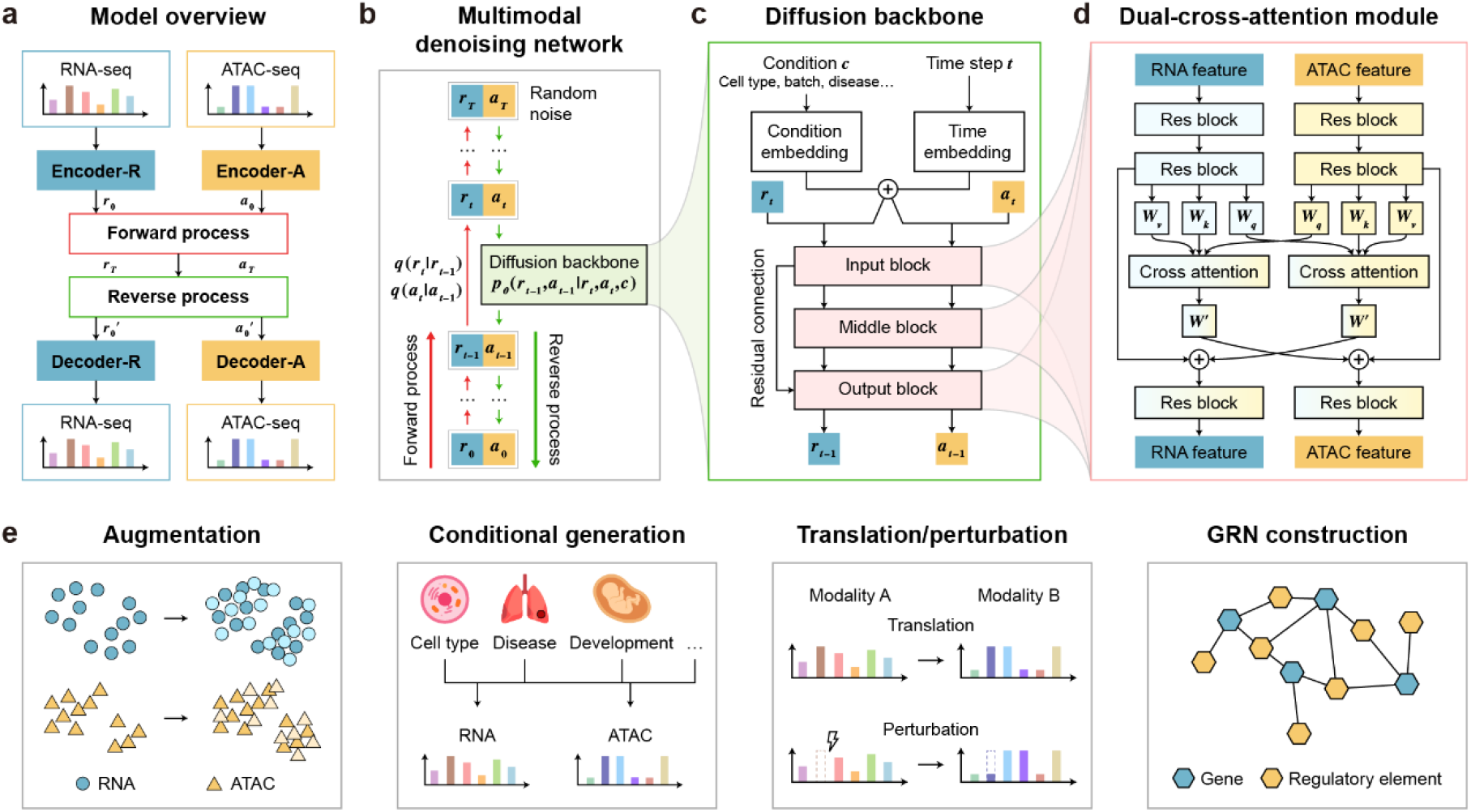
An interpretable deep generative framework for single cell multi-omics data analysis. (a) An overview of scDiffusion-X model. scDiffusion-X is a latent diffusion model that consists of encoders, decoders, coupled with the multimodal denoising network. (b) The multimodal denoising network. The forward process adds noise to the latent representation in each time step and the reverse process aims at removing the noise from a random generated noisy latent representation with the multimodal denoising network parameterized by *θ*. (c) The architecture of the multimodal denoising network from time t to t-1 where the condition and time step are used as additional inputs. (d) The Dual-Cross-Attention (DCA) module. This module uses a cross-attention mechanism to integrate information from different modalities. (e) scDiffusion-X model provides solutions to various downstream tasks.

During the training phase, the multimodal autoencoder first learns latent embeddings from paired multi-omics data. These embeddings, along with metadata as conditions, are then used to train the denoising network, which learns to reconstruct meaningful multi-omics signals from random noise. In the inference phase, the denoising network generates latent embeddings for different modalities from Gaussian noise, conditioned on user-specific inputs, which are then decoded back into their respective omics modalities by the multimodal autoencoder. We applied the scDiffusion-X model to multiple downstream tasks (Figure 1e), including multi-omics data argumentation, multi-omics data conditional generation, modality translation and perturbation, and gene regulatory network inference. By integrating the power of diffusion models, cross-attention mechanisms, and condition-aware learning, scDiffusion-X offers a novel and interpretable framework for generating and analyzing single-cell multi-omics data, providing new insights into cellular heterogeneity, regulatory interactions, and biological mechanisms.

### 2.2 scDiffusion-X Generates Realistic Single-cell Multi-omics Data

With hundreds of computational methods developed for various analytical tasks of single-cell multi-omics data^33-37^, fair and reliable benchmarking requires realistic data simulators that mimic the statistical properties and underlying distributions of real single-cell multi-omics data while retaining biological relevance^38,39^. In silico simulation methods that generate biologically realistic data with known ground truth have become essential for benchmarking computational tools.

To evaluate the ability of scDiffusion-X to generate realistic multi-omics data, we trained the model with two multiome datasets from the OpenProblem^40^ and 10x Genomics (PBMC10k) separately using cell type as condition labels and generated paired scRNA-seq and scATAC-seq data in proportion to the cell types in the original dataset. The goal was to assess whether scDiffusion-X could accurately learn the underlying distribution of real data while preserving key biological signals. For comparison, we also generated multi-omics data using other simulators, including MultiVI^15^, CFGen^21^, and scDesign3^24^, as well as two single-modal generators: scVI^16^ and peakVI^17^, which generate scRNA-seq and scATAC-seq data respectively. During the experiments, we found that the statistical model scDesign3 struggled with scalability, limiting its application to datasets with a large number of features and cells. To ensure a fair comparison, we filtered the features and cells in the original dataset, using the filtered dataset to train both scDesign3 and scDiffusion-X. Details on data processing and model configurations can be found in Methods.

We evaluated the performance of scDiffusion-X alongside baseline models using various metrics. The results demonstrated that scDiffusion-X generated more realistic multi-omics data compared to state-of-the-art methods. The UMAP visualization of data generated by scDiffusion-X exhibited high consistency to real data (Figure 2a-b, and Figure S1a-b), effectively preserving the global structure of real data, maintaining distinct cellular clusters, and capturing pseudotime information revealed from the scRNA-seq. Compared to baseline models, scDiffusion-X generated cells closely resemble real cells, while other methods, such as CFGen and MultiVI, exhibit distortions in cluster organization. Furthermore, quantitative evaluations across multiple metrics (see Methods), including Spearman Correlation Coefficient (SCC, which measure the degree of correlation between real and generated data), Maximum Mean Discrepancy (MMD, which measure the distance of real and generated data distributions), Local Inverse Simpson’s Index (LISI, which measure the batch effect of real and generated data distributions), and Area Under the Curve (AUC, which measure the overall similarity of biological landscape), were used to assess the performance for different methods with rankings visually indicated by color gradients, where lighter colors represent better performance and darker colors indicate lower rankings (Figure 2c, Figure S1c). Note that since AUC score quantifies how well an external Random Forest classifier can distinguish between real and generated data, lower AUC scores indicate greater similarity between generated and real cells. For scRNA modality, scDiffusion-X consistently achieves top performance across most metrics, indicating strong agreement with real data distributions. Notably, scDifussion-X improved the MMD and LISI by 33.3% and 15.5% compared to the second-best method. For scATAC modality, scDiffusion-X also demonstrates superior performance, ranking highest in SCC, MMD, LISI, and AUC scores, showing its effectiveness in capturing meaningful chromatin accessibility patterns. The AUC score achieved by scDiffusion-X was substantially improved from 0.846 to 0.575, compared to the best baseline, demonstrating the ability to generate realistic and indistinguishable chromatin accessibility profiles that closely mirror experimental data.

**Figure 2.**
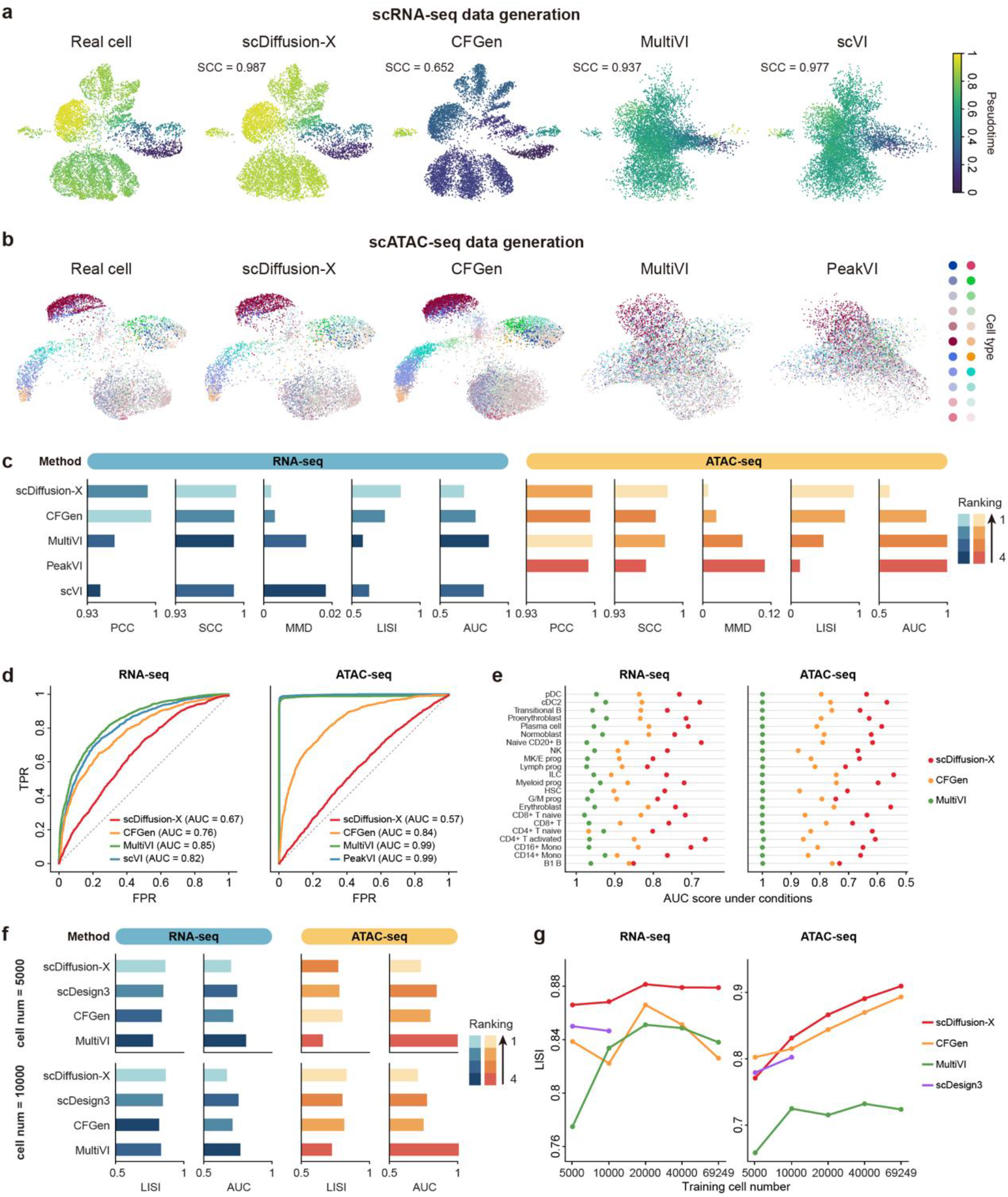
scDiffusion-X generates realistic single cell multiomics data. (a) The UMAP of generated scRNA-seq data by different methods on Openproblem dataset with pseudotime. (b) The UMAP of generated scATAC-seq data by different methods on Openproblem dataset. (c) Performance of different methods on Openproblem dataset under different evaluation metrics. (d) ROC curve of the external Random forest classification for the generated data and real data. AUC values closer to 0.5 indicate better generation quality. (e) Performance when generating under specific cell type condition. AUC values closer to 0.5 indicate better generation quality. (f) Performance of different methods on filtered Openproblem dataset under different evaluation metrics. (g) The model performance under different training data sizes.

In addition to global distribution preservation, we evaluated whether scDiffusion-X could generate realistic cell-type-specific multi-omics data. For comparison, we used CFGen and MultiVI as baseline models, we generated multiome profiles given specific cell types and assessed the AUC scores. As shown in Figure 2e, the cells generated by scDiffusion-X closely resembled real cells, achieving lower AUC scores than those generated by the baseline models. Specifically, the average AUC score of scRNA-modality improved from 0.864 to 0.747 and the average AUC score of scATAC-modality improved from 0.802 to 0.640 compared to the best baseline. To further validate the biological fidelity of the generated data, we analyzed the distribution of marker genes for each cell type against real cells. The Kullback-Leibler (KL) divergence score (Figure S1d) indicated that scDiffusion-X effectively retained key biological signals across different cell types. Additionally, the correlation patterns between the top differentially expressed genes and differential chromatin peaks in different cell types exhibit strong concordance in generated and real data (Figure S1e). These findings indicate that scDiffusion-X effectively preserves gene regulatory relationships across both modalities.

To explore the practical benefits of scDiffusion-X, we tested whether generated rare cell types could enhance downstream computational analyses, such as rare cell type identification. Specifically, we used cDC2 and plasma cells generated by scDiffusion-X to augment the original OpenProblem dataset and trained a Random Forest classifier for multi-class cell type classification. The results revealed that adding simulated rare cells significantly increased the F1 score of the rare cell type from 0% to more than 80% while maintaining high accuracy for other cell types (Figure S1f). These findings suggest that scDiffusion-X can serve as an effective data augmentation tool, enabling better representation of underrepresented cell populations in multi-omics studies.

A key advantage of scDiffusion-X over existing generative models is its scalability, particularly when handling large datasets with high-dimensional features. As shown in Figure 2f and Figure 2g, we randomly filtered the OpenProblem dataset into different sizes, filtered the features according to their variability (see Methods), and trained scDiffusion-X and the baseline models on these filtered datasets. We found that the statistical model scDesign3 failed to scale effectively to larger datasets with more than 10,000 cells, while scDiffusion-X still improves the performance as the dataset size increases, with the LISI score rising from 0.86 to 0.88 in RNA modality and from 0.77 to 0.90 in ATAC modality. Moreover, scDiffusion-X consistently achieved the best results in LISI score compared to other deep-learning-based models, emphasizing its robustness and adaptability in large-scale data generation. Overall, these findings demonstrate that scDiffusion-X is capable of generating realistic single-cell multi-omics data (e.g., multiome) by effectively preserving global data distributions, condition-specific signals, and gene regulatory relationships under specific conditions.

To make our model more useful, we trained the model on the latest scRNA-seq and scATAC-seq multi-omics dataset MiniAtlas^41^. This dataset contains more than 130,000 paired scRNA-seq and scATAC-seq data from 56 different cell types. scDiffusion-X again demonstrates superior performance on this large-scale dataset where the UMAP shows that the generated cells are ideally aligned with the real cell (Figure S2). We provide the pretrained model weight and training dataset on our project website (see Code availability). The user can use this pretrained model directly or finetune this base-model on their own data for user-specific task.

### 2.3 scDiffusion-X Enables Modality Translation and in-silicon Perturbation

The multi-omics sequencing technique continues to face experimental challenges, with costs much higher than those single-modality sequencing techniques^10,14,42^. Modality translation refers to generating one modality from another. It offers a powerful computational solution to overcome these limitations, effectively expanding the data from a single modality to multi-omics data^40^. Unlike existing multi-omics simulators, which fail to translate between omics modalities, scDiffusion-X uniquely enables direct cross-modality generation after model training. The model was trained with both modalities, as in the previous section, and did cross-modality generation by giving only one modality to generate the other from noise. Specifically, for each timestep t during the reverse process of the denoising diffusion model in the inference phase, we utilized the ground-truth noisy latent representation from the known modality in the forward process to replace the model-predicted noisy latent representation, generating the other modality from the standard Gaussian noise. The noisy latent representation served as a condition, guiding the model to generate the other modality through the multimodal denoising network with DCA (see Methods).

To assess the effectiveness of scDiffusion-X, we compared the scDiffusion-X model to the state-of-the-art modality translation model BABEL^32^ and evaluated the performance of both methods on the BABEL dataset. Following the original study, we clustered the scRNA-seq data in the dataset using the Leiden algorithm^43^, reserving one cluster (both scRNA-seq and its corresponding scATAC-seq) for validation and one for testing, while utilizing the remaining clusters for training. Additionally, we used the external GM12878 cell line with both scRNA-seq and scATAC-seq datasets for a completely out-of-distribution test. We trained the model in the training set and performed modality translation in the test set and the external dataset. As illustrated in Figures 3a-b, the translated scRNA-seq and scATAC-seq data closely resemble the real data in the UMAP visualization. The evaluation metrics demonstrated that scDiffusion-X outperformed BABEL in both RNA to ATAC and ATAC to RNA modality translation tasks. Specifically, scDiffusion-X achieved LISI scores of 0.55 and 0.67 for the translated scRNA-seq and scATAC-seq, compared to only 0.05 and 0.31 for BABEL. Besides, scDiffusion-X achieved MMD scores of 0.26 and 0.18 for translated scRNA-seq and scATAC-seq in the test cluster, which is 83% and 54% lower than that in BABEL (Figure 3c). These substantial improvements indicate that scDiffusion-X better preserves cell-type diversity and structure in the translated single-cell data. These results were further validated using the external dataset (Figure S3). Furthermore, the pseudo bulk profile based on average gene expression across the generated cells exhibited a better alignment (e.g., with a slope closer to 1) with the real data compared to BABEL (Figure S4). We also assessed the distribution of marker gene expression in the translated test set, revealing that the Kullback-Leibler divergence (KLD) scores of scDiffusion-X translated cells were lower than that of BABEL, indicating more effective reconstruction of biological signals (Figure 3d). scDiffusion-X reduced the KLD by 59.8% on average for the 10 marker genes provided from the original study^44^. It is seen that scDiffusion-X better captures the true distribution of marker genes, even if it is a multimodal distribution (e.g., ODC1), while BABEL can mostly learn an unimodal distribution. These results suggest that scDiffusion-X successfully captures and reconstructs the cross-modality relationships between RNA and chromatin accessibility signals, allowing for accurate modality translation. To further evaluate whether scDiffusion-X learned the functional relationships between modalities, we conducted a cross-modality perturbation experiment where we manually knocked out a specific gene (set to zero) and observed the resulting changes in the translated scATAC-seq. The rationale behind our experiment is that if the model has fully learned the interrelationship between the two modalities, the translated scATAC-seq should reflect the changes corresponding to a perturbed gene. To enhance the model’s sensitivity to these perturbations, we first trained it on a smaller dataset with highly variable genes and peaks. Next, we compared the translated scATAC-seq with the results from a real perturbation experiment^45^. Specifically, we calculated the differential peaks between the perturbed and control groups in the actual experiment, merged these groups into an average bulk cell, and determined the change direction of the top differential peaks after the perturbation. We applied the same procedure to the diffusion-translated control and perturbed peaks and assessed the accuracy of the change direction relative to the real perturbation experiments. We also used the MultiVI model with the same settings as a comparison. The results indicated that scDiffusion-X successfully predicted over 80% of the change directions among the top 20 to 50 differential peaks in the perturbation of ZAP70, CD3E, and CD4 (Figure 3e), demonstrating its capability to capture perturbation-induced regulatory changes and adjust the corresponding modality accordingly. The accuracy declined to around 75% for the top 100 differential peaks, exhibiting weaker perturbation effects. We also performed cross-modality perturbation experiment on a real multiome perturbation dataset^46^, and the results showed that scDiffusion-X outperformed the MultiVI model in the prediction of change direction after perturbation (Figure S6). These results highlight the ability of scDiffusion-X to accurately translate missing omics modalities, reconstruct biological signals, and capture cross-modality perturbation effects.

**Figure 3.**
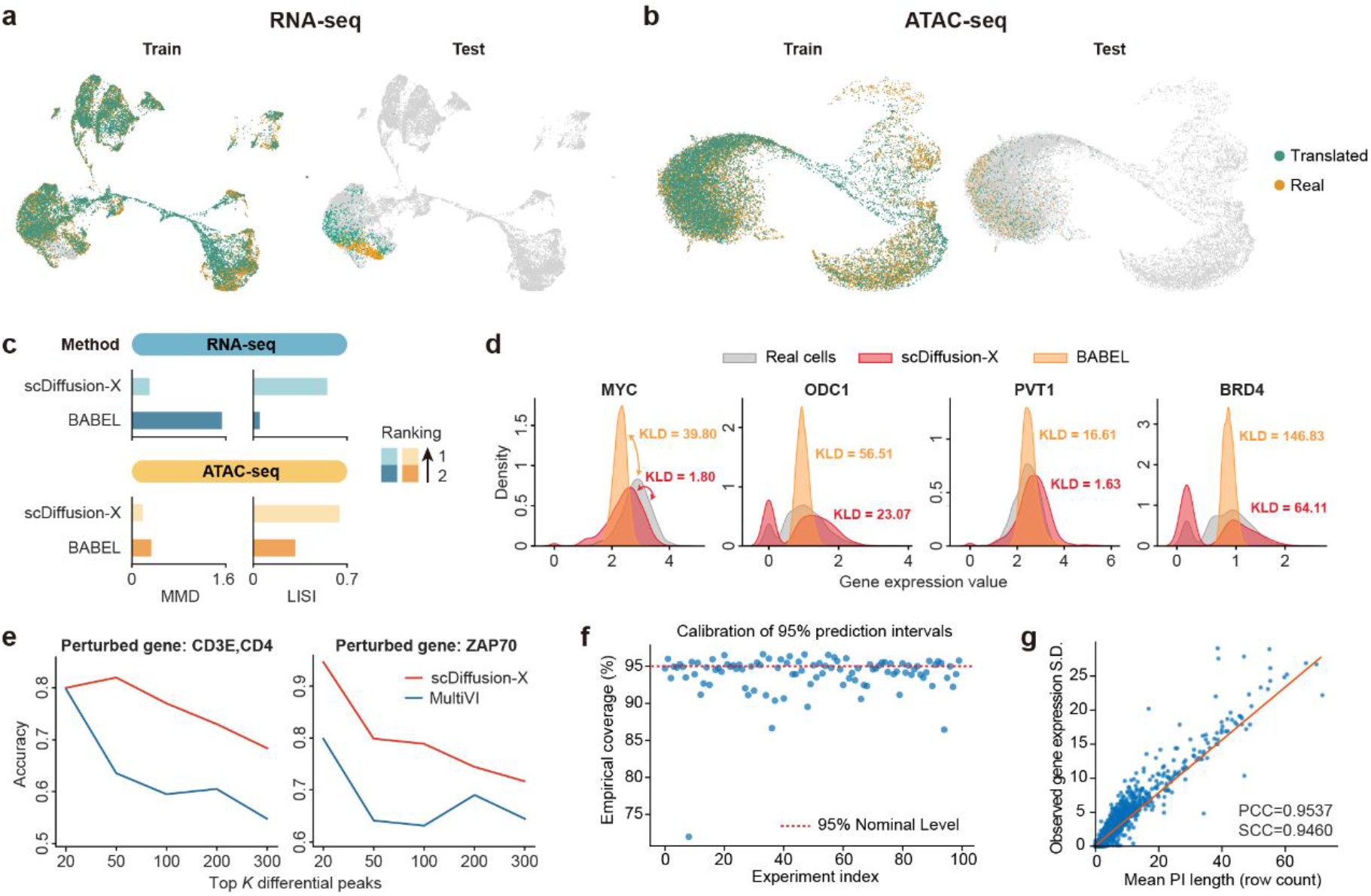
scDiffusion-X enables modality translation. (a) The UMAP of translated scRNA-seq and real scRNA-seq data from test set and training set. (b) The UMAP of translated scATAC-seq and real scATAC-seq data from test set and training set. (c) Performance of scDiffusion-X and BABEL on test set under different evaluation metrics. (d) The comparison of gene expression distribution of test set marker genes between real data and translated data by two methods. (e) The change direction’s accuracy of top differential gene/peak after perturbation by scDiffusion-X and MultiVI. (f) The proportion of real gene expression falls within the 95% prediction interval in each of the 100 times experiments. (g) The relationship between prediction interval (PI) length and observed gene expression S.D..

To evaluate the model’s ability to capture data uncertainty from the observed single-cell multiomics data, we constructed prediction intervals (PI) for uncertainty quantification by multiple runs of modality translation from scATAC-seq to scRNA-seq with independent random noise (see Methods). As shown in Figure 3f, we repeat this process 100 times, and the model demonstrated excellent calibration by demonstrating an average coverage proportion of 93.94% across genes, closely matching the nomimal 95% level, indicating strong calibration. We further examined whether the uncertainty captured by scDifussion-X truly reflects the observed variation of expression level in real data across genes. To do this, we first calculated the standard deviation (S.D.) of each gene in the real test set data, and then calculated the average length of PIs over 100 experiments. Ideally, genes with higher variance are expected to exhibit a wider prediction interval. We quantified this relationship using the Pearson correlation coefficient (PCC) and Spearman correlation coefficient (SCC). The analysis revealed an average PCC of 0.9537 and SCC of 0.9460 between PI length and gene expression S.D.(Figure 3g), demonstrating that the model effectively captures the inherent gene-specific uncertainty. The expression distributions of the top expressed genes across 100 generated profiles also closely matched the real distributions (Figure S5), further confirming that the model learned an full generative distribution, providing richer information than point estimates.

### 2.4 scDiffusion-X Uncovers Gene Regulation Mechanisms from DCA

A fundamental challenge in single-cell multi-omics analysis is deciphering the intricate relationships between different molecular layers to uncover the underlying gene regulatory mechanisms^8,47^. To investigate this and provide biological insights for the generation process, we proposed a gradient-based interpretability approach to analyze how different omics modalities interact in the DCA. Specifically, the DCA encodes the interaction information within the cross-attention matrix, each element in the attention matrix quantifies the degree to which one modality attends to another. We computed the gradient of the latent elements in the cross-attention matrix for all input genes or peaks. The magnitude of these gradients indicates the importance of each gene or peak relative to the elements (Figure 4d), thereby establishing a regulatory interaction map between genes and chromatin accessibility regions (see Methods). We used the model trained on the OpenProblem dataset to analyze these interactions between modalities in this experiment.

**Figure 4.**
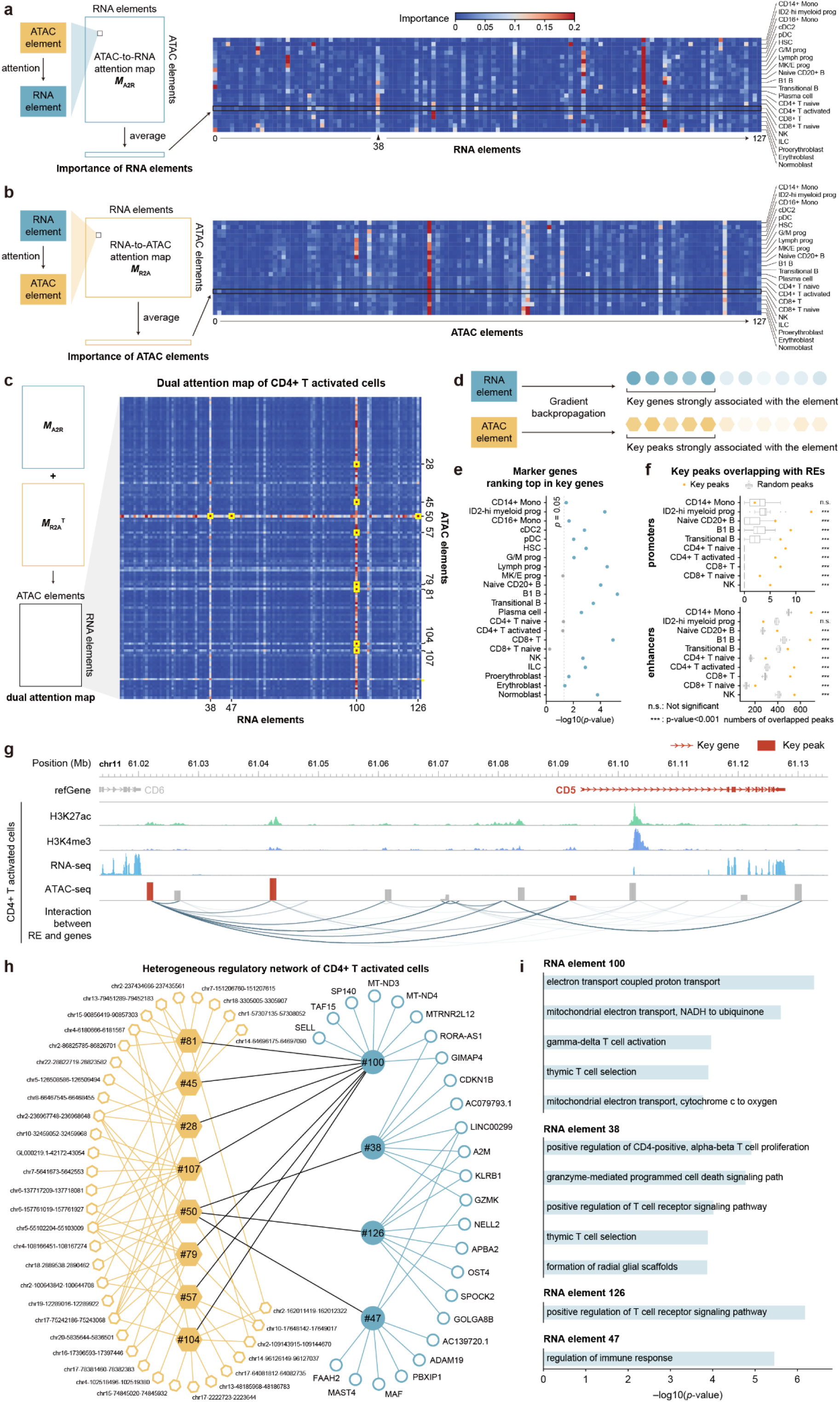
(a) Aggregated attention map of each RNA element to ATAC elements in different cell types in the second DCA. (b) Aggregated attention map of each ATAC element to RNA elements in different cell types in the second DCA. (c) Dual attention map of CD4+ T activated cell and its construction process. (d) An illustration of gradient-based interpretability approach. (e) The significance test of marker genes’ gradient magnitudes rank. (f) The number of peaks overlaps with the promoter or enhancer regions. The dot is the number of diffusion-selected key peaks that overlap with the promoter (upper figure) or enhancer (lower figure). Boxplot is the range of randomly selected peak overlap with the promoter (upper figure) or enhancer (lower figure). Here we randomly selected 20 times and calculated the one-sample t-test with the diffusion overlapped number. (g) A showcase of key genes and the key peaks in CD4+ T activated cells selected by scDiffusion-X. The key peaks are located in the potential enhancer position and promoter position of the key gene CD5. (h) A showcase of the CD4+ T activated cell heterogeneous gene regulatory network inferred by scDiffusion-X. This network shows the relationship of key genes – key RNA elements – key ATAC elements – key peaks. (i) The GO terms obtained from the key genes of each RNA element.

The gradient-based interpretability approach and DCA can be used to identify the cell-type-specific highly-attended genes or peaks. We first identified the genes that were most attended to by peaks within each cell type. There are three DCA module in the scDiffusion-X, we take the second DCA module as an example. We averaged the ATAC-to-RNA attention map across cells within each cell type and then averaged it out at the ATAC axis, resulting in a *d*_*g*_-dimensional vector where *d*_*g*_ is the feature dimension in this DCA, representing the importance of each RNA element (Figure 4a). For the RNA elements with high attention scores, we employed the gradient-based interpretability approach to each of them to determine the strongly associated genes. After that, we chose the 100 genes with the largest gradient values and analyzed the potential Gene Ontology (GO) terms associated with these genes. We found that the GO terms of these genes are highly related to the cell type of the target RNA element. For example, the genes associated with the 38th element of CD4+ T cells in Figure 4a correspond to several immune-related GO terms (Figure S7a), such as negative thymic T cell selection (p-value=2.09×10 ^-5^) and the gamma-delta T cell receptor complex (p-value=1.05×10 ^-4^). We can also observe the attention scores of the 38th element in other immune cells are higher than other cell types, indicating its cell type specific for immunity (p-value=2.45×10 ^-5^, one-sided t-test). Similar enrichment patterns were observed for other gene sets, revealing key biological processes associated with specific attention-derived latent elements (Figure S7b-d). The same procedure was conducted to obtain the cell type specific importance of ATAC elements (Figure 4b) and elements in other DCAs (Figure S8a-b). Additional detailed analyses of peaks associated with specific ATAC elements are available in Figure S7e-f.

We calculated the gradient magnitudes of each gene for the top-5 high attention score RNA elements in each cell type and obtained the magnitudes rank of their marker genes given from the original study. We then used the rank sum test to assess whether the gradient magnitudes of marker genes for each cell type ranked above those of all other genes. The results revealed that the marker genes in 18 of the 22 cell types ranked significantly higher than others, indicating that the gradient-based interpretability approach effectively selects functionally relevant genes (Figure 4e).

The gradient-based interpretability approach and DCA can also be used to further investigate the regulatory relationships between different modalities. We examined the interactions between RNA and ATAC, represented through a dual attention map 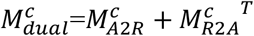 (Figure 4c). Here, 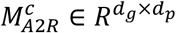denotes the attention of genes towards peaks, while 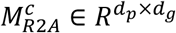 captures the attention of peaks toward genes. Latent elements with high attention weights indicate that the RNA and ATAC are mutually attentive to each other (Figure 4c, Figure S8c-d). By applying gradient backpropagation to these latent elements with high attention scores, we identified a set of key genes and peaks that are likely to exhibit a regulatory relationship.

To validate these inferred regulatory relationships, we compared our identified key peaks and key genes against ENCODE H3K27ac data^48,49^ and chromatin loop data from the HiChIP database^50^. Our findings showed that the majority of key peaks are located within enhancer or promoter regions or directly overlap with the associated key genes. A one-sample t-test revealed that our key peaks have significantly higher chances of overlapping with enhancer markers and transcription start sites (TSS) of genes compared to randomly selected genes and peaks in most cell types (Figure 4f). We further investigated the more specific potential regulatory relationships of these key genes and peaks. We found the locations of these key genes on the chromosomes and then focused on the key peaks around these key genes to see if they were located on the potential regulatory element positions of these key genes. For instance, as illustrated in Figure 4g, three key peaks near key gene CD5 function were identified as potential promoters and enhancers regulating CD5 expression. Similar regulatory patterns were observed across multiple DCAs and across different cell types (Figure S9a-b).

To provide a global view of gene regulation, we constructed a heterogeneous gene regulatory network for each cell type that connects genes, peaks, RNA elements, and ATAC elements. This approach provides a novel way of interpreting gene regulation by interpreting the DCA in the latent space (see Methods). Unlike traditional gene regulatory network (GRN) studies, which primarily rely on co-expression patterns^51,52^, chromatin interaction maps^53,54^, or predefined enhancer-promoter links^55,56^, the DCA in our approach enables genes and peaks to interact in low-dimensional latent spaces, allowing the model to learn regulatory relationships directly from data rather than relying on manually curated annotations. Here, the term “heterogeneous” means the nodes and edges in the network have different types and attributes. Specifically, the resulting heterogeneous gene regulatory network integrates four types of regulatory elements (nodes in the regulatory graph, including gene, RNA element, ATAC element, peak) and defines connections based on the strength of learned interactions (edges in the regulatory graph). As demonstrated in Figure 4h (and also Figure S10a-b), this heterogeneous network has four RNA elements and eight ATAC elements, each of them linking 8 top-associated genes/peaks. It’s noted that some elements (e.g., the 100th RNA element) have large degrees, showing their importance in the regulatory network. Some key real peaks/genes, like gene GZMK and peak chr17-75242186-75243068, are linked to multiple latent elements, suggesting their potential biological function to this specific cell type. We found that the GO terms associated with these key genes are highly relevant to the underlying cellular context. For example, for CD4+ T activated cells, the GO terms of the real gene linked to the 38th RNA element included GO terms like *regulation of CD4-positive, alpha-beta T cell proliferation*, and *regulation of T cell receptor signaling pathway* (Figure 4i, see other examples in Figure S10c).

We then linked the gene and peak through the heterogeneous network. Specifically, if a peak and a gene can be linked through an ATAC element and an RNA element, we select these peak-gene pairs as a potential regulatory relationship. We restricted these pairs to intra-chromosomal interactions and further filtered out pairs where the peak–gene distance exceeded 120 kbp. Using the heterogeneous network of CD4+ activated T cells as an example, our approach identified 366 gene-peak pairs. We used the HiChIP loop data from HiChIPdb^50^ as the ground truth for the regulatory relationship to see how many selected pairs can be validated by HiChIP loops. We compared our results with several baselines, including the state-of-the-art GRN method, such as SCENIC+^57^ (see Methods). The results showed that the peak-gene pairs selected by scDiffusion-X reach a 0.64 precision score (**Table S2**) and surpass all baseline methods. This indicated that the heterogeneous GRN inferred by scDiffusion-X can capture biological associations between genes and chromosomal regions, providing insights into potential regulatory relationships. Overall, this heterogeneous gene regulatory network offers a data-driven approach to studying gene regulation, providing biological interpretability for the generation process of multi-omics data.

### 2.5 Evaluating the Contribution of the DCA

A key challenge in multi-omics data integration is effectively capturing the relationships between different modalities. A common approach is to concatenate multi-modalities in an early or late stage. However, this simple concatenation treats modalities as independent features and fails to model their complex dependencies explicitly. In contrast, the DCA in scDiffusion-X provides a more interpretable way to integrate multi-omics data by explicitly modeling interactions between different modalities at each step of the denoising process. To assess its significance and the richness of the information it captures, we conducted a series of ablation studies using the OpenProblem dataset. First, we varied the number of DCAs by one, three, and five to evaluate their impact on model performance. Then, for the model with three DCAs, we substituted the DCA with the fully connected linear layer that concatenated the latent representations of different modalities, allowing them to interact through the linear layer. As shown in Figure 5a, model performance improved progressively with an increasing number of DCAs, highlighting the beneficial role of this module. In contrast, the linear layer approach yielded suboptimal performance compared to the cross-attention module, indicating that the latter captures the interactions between different modalities more effectively. However, the addition of more DCAs increases computational costs. Balancing model performance with computational efficiency, we ultimately use three DCAs as the default configuration.

**Figure 5.**
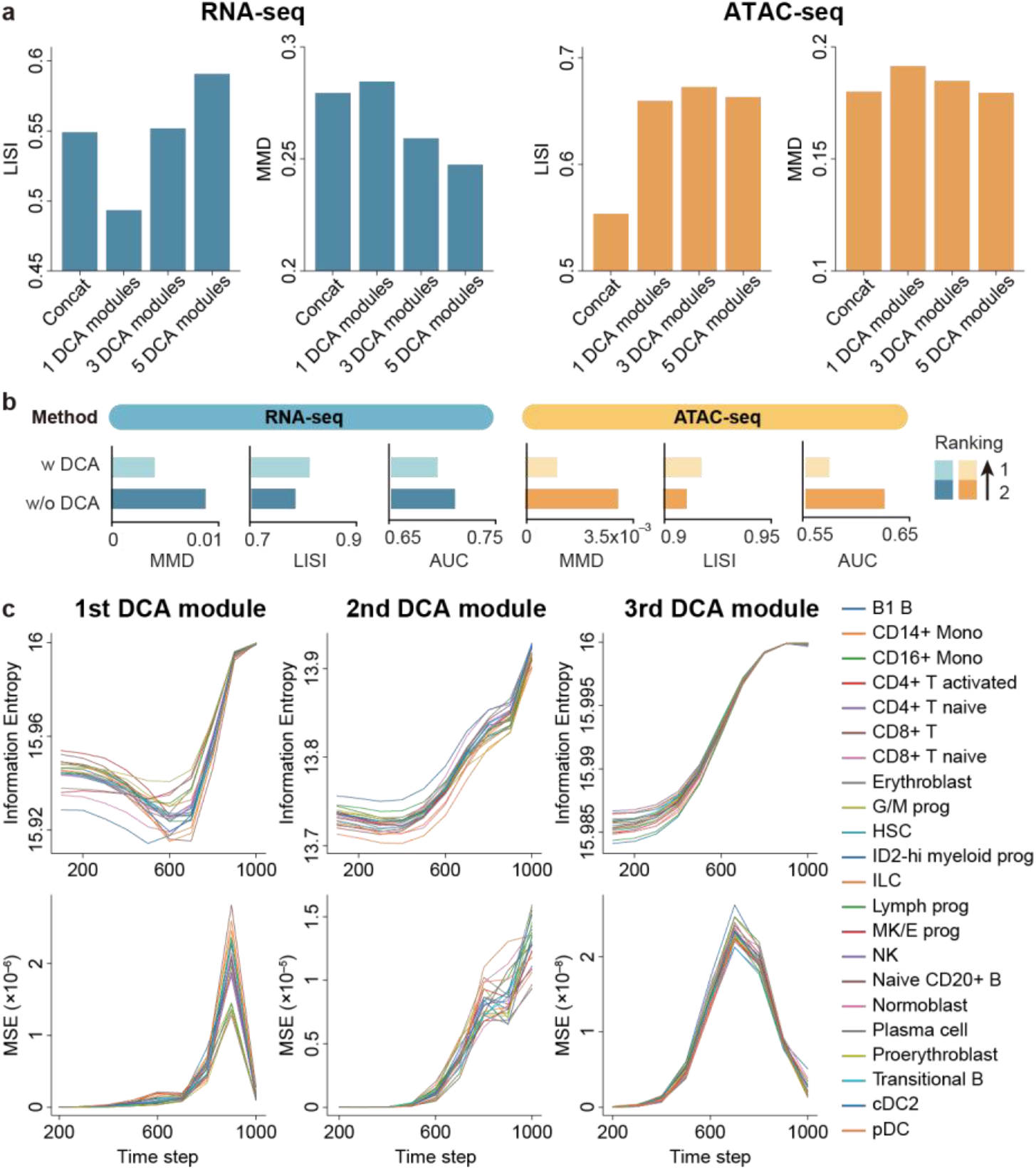
Investigating the effectiveness of the DCA. (a) Ablation study for DCA. We set the number of DCAs to 1, 3, and 5 to train the scDiffusion-X model. We also used 3 linear layers to replace 3 DCAs to verify the effectiveness. (b) The performance of scDiffusion-X with(w) or without(w/o) the DCA. (c) Ablation study for information richness of ATAC to RNA attention maps in different DCAs and different time steps.

We then removed the DCA to investigate whether the model can leverage information from an alternative modality to facilitate the generation of the current modality. To minimize the variables introduced by the additional information, we trained the model without cell type conditions. As illustrated in Figure 5b, the model trained with the DCA exhibited superior performance on MMD, LISI, and AUC metrics for both scRNA-seq and scATAC-seq data, compared to the model without the DCA. This indicates that the model benefits from the information extracted from the other modality by DCA when generating the current modality.

The model incorporates multiple DCAs and numerous time steps in the diffusion process. The information richness in different DCAs and at various time steps varies considerably. To identify the most informative DCAs and time steps, we calculated the information entropy (IE) and mean square error (MSE) of the attention maps across different modules and steps using the model trained on the Openproblem dataset. The IE quantifies the information richness within a specific attention map, while the MSE assesses the change in information compared to the attention map from 100-time steps earlier. As illustrated in Figure 5c (ATAC to RNA attention map) and Figure S11 (RNA to ATAC attention map), the IE increases across all three DCAs and all cell types as time progresses, indicating that the generated representations become more realistic and information-rich. The MSE exhibits similar trends in the first and second modules, reaching the maximum at the 900th time steps and 1000th time steps, whereas the third module demonstrates significantly lower values, suggesting that the first and second modules retain more informative signals for learning. Based on the insights gained from these two metrics, we selected the attention map from the second attention module at the 1000th time step and the first attention module at the 900th time step to elucidate the gene-peak regulatory mechanism. These results highlight the importance of the DCA over simple concatenation, demonstrating that the DCA not only enhances model performance but also improves interpretability by explicitly modeling the interactions between modalities.

## 3. Discussion

In this study, we presented scDiffusion-X, a latent diffusion model designed for single-cell multi-omics data. By leveraging a denoising diffusion framework and an innovative Dual Cross-Attention (DCA) module, scDiffusion-X enables high-fidelity data generation, modality translation, and gene regulation inference. Through comprehensive benchmarking, we demonstrated that scDiffusion-X achieves state-of-the-art performance in generating realistic single-cell multi-omics data, preserving cell-type specificity, and improving biological interpretability. A key advancement of scDiffusion-X is its ability to learn cross-modality dependencies adaptively through the DCA, offering a more flexible and interpretable alternative to traditional multi-omics integration methods. Furthermore, our gradient-based interpretability approach enables the DCA to construct a cell-type specific heterogeneous gene regulatory network that reveals comprehensive gene regulatory relationships in a data-driven manner.

Despite its strong empirical performance, there are several limitations that require further development and expansion. First, the current scDiffusion-X only supports the joint modeling of two modalities, such as multiome. Extending the architecture to accommodate other types of multi-omics data beyond two modalities would further enhance its applicability to complex biological systems. Second, scDiffusion-X does not yet incorporate spatial or temporal information, which are critical for understanding tissue organization and dynamic cellular processes. Integrating spatial omics data, such as spatial transcriptomics and proteomics, or time-series single-cell data could enable the modeling of spatiotemporal gene regulation, tissue development, and disease progression. Third, the growing availability of large-scale multi-omics datasets, such as the MiniAtlas dataset^41^ and scMMO-atlas^58^, provides an opportunity to pretrain scDiffusion-X on diverse datasets, covering a wide range of cell types and biological conditions. A pretrained foundation model for multi-omics generation could enable users to generate high-quality multi-omics data without the need for extensive model retraining, significantly reducing computational costs and making multi-omics simulation more accessible. We have released a model pretrained on the MiniAtlas dataset to enhance the usability of scDiffusion-X, and will further scale the data and model in the future. Finally, we note that many cross-modality methods, including ours, often focus on capturing the shared information between modalities, an assumption that has proven effective in domains such as language and vision. However, in biology systems, each modality often reflects disinct and complementary aspects of cellular states. Leveraging both shared and modality-specific signals could provide a more comprehensive understanding of cellular complexity. Future work should explore strategies that balance these two perspectives to maximize biological insight.

In summary, scDiffusion-X provides a scalable, interpretable, and biologically grounded framework for single-cell multi-omics analysis. As single-cell technologies continue to evolve, scDiffusion-X has the potential to become a versatile tool for advancing our understanding of cellular heterogeneity, gene regulation, and disease mechanisms.

## 4. Methods

### 4.1 Vanilla Diffusion Probabilistic Models

Diffusion Models are probabilistic models designed to learn a data distribution *p*(*x*) by progressively denoising a variable sampled from a standard Gaussian distribution^59,60^. Diffusion Models consist of two main stages: a forward process and a reverse process. The forward process progressively corrupts observed data by incrementally adding Gaussian noise over a series of T steps until it conforms to a standard normal distribution. Conversely, the reverse process learns to iteratively denoise the data by training a parameterized model (e.g., neural network), which is structured as a Markov chain of T steps. Define *x*_0_ as a sample from *p*(*x*), and *x*_T_ as the corrupted sample that fits standard Gaussian distribution. The forward noise-adding process is described by the Markov forward process:

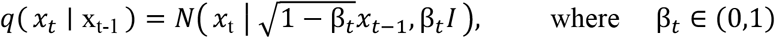

Where *t* ∈ [1, *T*] , *I* stands for standard Gaussian noise. *β*_*t*_ is a pre-defined variance schedule sequence that increases with time step.

The reverse process learns to reverse the forward process to recover the original data distribution. This can be simplified as training a denoising network Ө to fit *p*_Ө_ (*x*_*t*−1_ ∣ *x*_*t*_)for all given *t* and *x*_*t*_. Thus, the reverse process is formulated as:

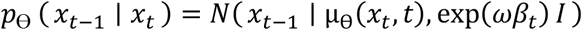

Where ω in the variance is an adjustable weight, and the *μ*_Ө_(*x*_*t*_, *t*)is the Gaussian mean value predicted by denoising network Ө. The Diffusion Model initiates with Gaussian noise and progressively denoises it over T steps to ultimately recover the sample *x*_0_:

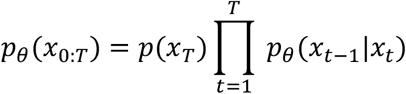

### 4.2 Multimodal Autoencoder for Embedding and Reconstructing Multiome Data

To improve the scalability and efficiency of the generative model and reduce the noise in the single-cell data, we adapted the latent diffusion model^61^ in scDiffusion-X. We used a multimodal autoencoder to embed the different modality data into low-dimensional latent representations and used these latent representations to train the denoising network.

The multimodal autoencoder consists of two separated autoencoders based on the Multilayer Perceptron (MLP) network. The detailed architecture and hyperparameters of the autoencoder can be found in Figure S12a. We referred to the architecture in CFGen and scVI to train the autoencoder, which assumes that gene expression values follow a negative binomial distribution, and chromatin accessibility conforms to a Bernoulli distribution. The training target is to maximize the log-likelihood of the data. Here, we take the scRNA-seq and the scATAC-seq as an example. For scRNA-seq data, the RNA autoencoder first encodes the scRNA-seq profile *x*_*R*_ into latent representation *r* , and then use the decoder to get the likelihood parameters, which use a negative binomial distribution *x*_*R*_∼*NB*(*μ, θ*)to fit the scRNA-seq and can be inferred as:

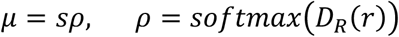

Where *D*_*R*_ is the RNA decoder, *s* is the size factor which is the total counts of scRNA-seq data, and the inverse dispersion parameter *θ* is a learnable parameter for each gene. For scATAC-seq, the model uses the Bernoulli distribution *x*_*A*_∼*Bern*(*π*)to fit the scATAC-seq since the scATAC-seq are all binary, and the parameters *π* can be inferred as:

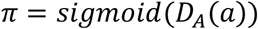

Where *D*_*A*_ is the ATAC decoder, *a* is the latent representation of scATAC-seq. These log-likelihood losses enable both of the autoencoders to accurately reconstruct the input data from a low dimensional feature space, reducing the complexity of the denoising network.

### 4.3 Multimodal Denoising Network for Data Generation

The multimodal denoising network is to generate new latent embeddings of different modalities from standard Gaussian noise. Following the standard training approach for latent diffusion models, we applied the diffusion process to the latent embedding derived from the multimodal autoencoder and generated noisy data to train the denoising network.

Specifically, given paired data (*x*_*R*_, *x*_*A*_)from scRNA-seq and scATAC-seq, we first used the trained multimodal autoencoder to get the latent embedding *r*_0_ and *a*_0_, and then applied the forward diffusion process to obtain the noisy embedding *r*_*t*_ and *a*_*t*_ (Figure 1b). The forward processes for different modalities can be considered independently, which are defined as below:

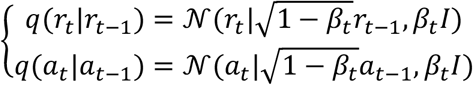

Where *t* is the time step, *I* stands for identity matrix, and *β*_*t*_ is a coefficient that varies with the time step:

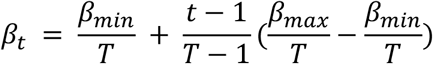

where *β*_*min*_ and *β*_*max*_ are two parameters that control the scale of *β*_*t*_ in the diffusion process. Here we set the *β*_*min*_ to 0.0001 and *β*_*max*_ to 0.02. During the reverse process, considering that there is a complex relationship between different modalities, we designed a multimodal denoising network to take both modalities as input and fully interact the information of these two modalities through the DCA to improve the generation quality. Specifically, considering noisy representations *r*_*t*_ and *a*_*t*_, the reverse process can be written as:

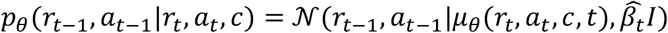

Where 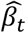 is the coefficients that vary with time step, 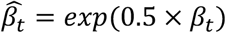, *c* is the input condition, and the *μ*_*θ*_ stands for the multimodal denoising network with learnable parameters *θ*.

As shown in Figure 1c-d, the multimodal denoising network consists of two single-modal denoising networks for different modalities generation. Each of them is in the typical U-Net form, consisting of Input layers, Middle layers, and Output layers, which are all connected with residual connection to avoid loss of information wherever possible. In the meantime, these two single-modal networks are coupled with the Dual-Cross-Attention module, which fully interacts the information of these two modalities and reinforces the generation process. It takes the latent representation of scRNA-seq and scATAC-seq and outputs the predicted noise to reverse the noisy data. The condition information is embedded and added to the time embedding to control the generation process. The detailed architecture and hyperparameters of the multimodal denoising network can be found in Figure S12b-e.

The Res block is the basic component of different layers. It is a residual connected layer that combines the time/condition embedding and the latent feature. Assume the input feature is *r*_*in*_ ∈ *R*^*n*^ (n is the feature dimension), the time embedding is *t*_*emb*_ ∈ *R*^*n*^ and the condition embedding is *c*_*emb*_ ∈ *R*^*n*^. The Res block can be formulated as:

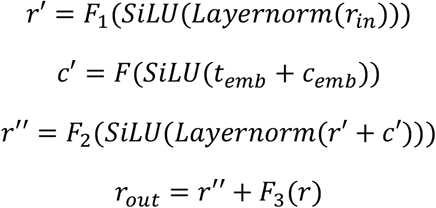

where *r*_*out*_ is the output feature and the *F*_1_, *F*_2_, and *F*_3_ are the linear layers. *SiLU* is the activation function, which can be defined as:

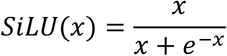

where *x* is the input. There are 3 Res blocks in the Input and Output block, two before the DCA and one after the DCA and the residual connection (Figure 1d). For the middle block, we remove the two Res blocks before the DCA for computational efficiency. The input feature size for three Res blocks in the Input block is 512, 256, and 128. For Res blocks in the Output block, it is 128, 256, and 512. For the middle block, the size is 128.

### 4.4 Dual-Cross-Attention Module for aligning RNA and ATAC modalities

To synthesize the information embedded within the two modalities, we developed the Dual-Cross-Attention module, which employs a cross-attention mechanism to assess the information one modality attends to another (Figure 1d). The foundamental assumption is that paired RNA-seq and ATAC-seq profiles obtained through multiome data contain intrinsic relationships and therefore provide mutual information gain. Our Dual-Cross-Attention module introduces a new mechanism to construct the Query (Q), Key (K), and Value (V) specifically for each modality, allowing both modalities to interact without concatenating them while preserving their original dimensionality.

Specifically, assume that for each cell from a multi-omics profile, scRNA-seq is represented by an n-dimensional feature vector R ∈ *R*^*n*×1^ , and scATAC-seq is represented by an m-dimensional feature vector A∈ *R*^*m*×1^. We first create Q, K, and V ∈ *R*^*n*×*d*^ for both modalities. For RNA-seq, we used a trainable matrix W_Q,R_ ∈ *R*^1×*d*^ to obtain the Q_R_ by Q_R_ = RW_Q,R_ ∈ *R*^*n*×*d*^, and used trainable matrix W_K,R_ ∈ *R*^1×*d*^ and W_V,R_ ∈ *R*^1×*d*^ to obtain the K_R_ and V_R_ by K_R_ =RW_K,R_ ∈ *R*^*n*×*d*^ and V_R_ = RW_V,R_ ∈ *R*^*n*×*d*^. For ATAC-seq, we used a similar process to obtain Q_A_, K_A_, and V_A_. Then the cross-attention is calculated in both directions as defined below:

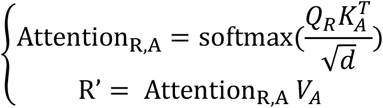

Where scRNA-seq borrows information from scATAC-seq. And:

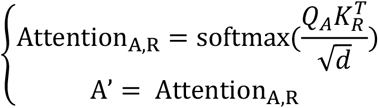

Where scATAC-seq borrows information from scRNA-seq. Note that the R′ ∈ *R*^*n*×*d*^ and A′ ∈ *R*^*m*×*d*^. Last, the outputs are mapped to the original dimension:

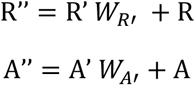

where W_R’_∈ *R*^*d*×1^ and W_A’_∈ *R*^*d*×1^ is the trainable matrix, R′′ ∈ *R*^*n*×1^ and *A*′′ ∈ *R*^*m*×1^. The +R or +A is a residual connection design in the network. There are totally three DAC modules in the scDiffusion-X model. One in the Input block, one in the middle block, and the other in the Output block. The feature dimensions in these three DCA are 256, 128, and 256.

### 4.5 Gradient-based Interpretability Approach and Heterogeneous Gene Regulatory Network Construction

To extract the corresponding genes and peaks to the RNA elements and ATAC elements in the DCA, we introduced the gradient-based interpretability approach. Specifically, for the attention map *M*^*c*^ given by one of the model’s intermediate DCAs, we use the backward() function in the torch library to calculate the gradient backpropagation of the latent elements of interest (usually the ones with high attention scores) in this attention map. After that, each input gene and peak will have a gradient to this attention map:

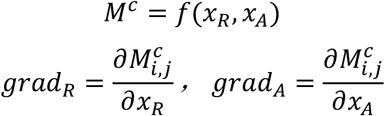

where *grad*_*R*_ and *grad*_*A*_ are the gradients of the input gene and peak, *f* stands for the part of the model before this DCA, and *i, j* is the position of the interested element in the attention map. The larger the magnitude of the gradient, the greater the effect of this gene/peak on this element on the attention map. In our experiments, we selected the top 100 genes/peaks with the largest gradient values as the key genes/peaks corresponding to the gene/ATAC elements. The heterogeneous gene regulatory network is constructed based on the dual attention map and the above gradient-based interpretability approach. Specifically, for a given cell type (or other conditions), we calculate the dual attention map 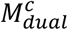 (Figure 4c) and select the RNA and ATAC elements with high attention scores. We choose the ten latent elements with the highest attention values and transform them into a bipartite graph given both the RNA and ATAC elements. We then use the gradient-based interpretability approach to obtain the genes and peaks corresponding to these elements, linking RNA elements back to genes and ATAC elements to peaks. The constructed heterogeneous gene regulatory network establishes a biologically meaningful mapping that provides insights into gene regulation at a fine-grained level. Note that for a clearer presentation, the heterogeneous gene regulatory network in Figure 4d only shows the top 10 associated genes or peaks for each element.

### 4.6 Uncertainty quantification and prediction interval

To evaluate the model’s ability to capture data uncertainty from the observed single-cell multiomics data, we constructed per-cell prediction intervals (PIs) for uncertainty quantification given a user-specific significance level *α*. The procedure was designed to assess the proportion of true gene expression values captured by the model’s generated distributions. Given M cells randomly selected from the test set, an ensemble of N=100 scRNA-seq profiles were generated by multiple runs of translation from scATAC-seq to scRNA-seq with independent random noise. This process yielded a predictive distribution for the expression of each gene. Note that the analysis was restricted to genes with non-zero expression in the ground-truth cell. For each of these expressed genes, a (1-*α*)% prediction interval (PI) was constructed by computing the *α*/2 -th and (1-*α*/2)-th percentiles from its N generated expression values, where *α* = 0.05 as default. The per-cell coverage was then calculated as the proportion of these expressed genes for which the ground-truth expression value fell within its corresponding PI. This procedure yields a single coverage score for each of the M cells tested. A well-calibrated model is expected to produce a distribution of per-cell coverage scores centered near the nominal (1-*α*)% level, indicating that the uncertainty estimates are reliable across the entire gene expression profile.

### 4.7 Baseline GRN method

To ensure robustness in GRN inference, we compared scDiffusion-X with several baseline strategies and established GRN inference methods:

#### Random pair

randomly selects peak–gene pairs within 120 kbp from the same chromosome. We randomly selected 366 pairs to match the number identified by scDiffusion-X.

#### Upregulated genes + nearest peak

selects the top upregulated genes in the CD4+ T cells and pairs each with its nearest peak. We selected the top 366 upregulated genes and their nearest peaks to match the number identified by scDiffusion-X.

#### Regulatory element (RE) + nearest gene

randomly samples regulatory elements from ENCODE H3K27ac data^48,49^ and assigns their nearest genes. We randomly selected 366 regulatory elements and their nearest peaks to match the number identified by scDiffusion-X.

#### MultiVI + correlation

denoises RNA-seq and ATAC-seq data using MultiVI^15^, and then calculates the Pearson correlation between gene expression and the openness of each chromatin region within a 120 kbp range around it. We selected the top 366 peak–gene pairs with the highest Pearson correlations within 120 kbp.

#### SCENIC+^57^

a classical GRN inference method, can obtain the importance scores for gene-peak pairs. The SCENIC+ identified 3642 possible pairs in CD4+ T cells, from which we selected the top 366 peak–gene pairs ranked by importance scores.

### 4.8 Dataset and Preprocessing

We used 4 datasets in our experiments, the Openproblem dataset^40^, PBMC10k dataset, the BABEL dataset^32^, and the MiniAtlas dataset^41^.

#### The Openproblem dataset

originated from the 2021 NIPS competition (Open Problems in Single-cell Analysis). It contains 69,249 cells from 22 different cell types. We filtered the gene that expressed in less than 10 cells and ATAC that opened in less than 2% of the cells, which remained 13,431 genes and 36,553 ATAC regions (peaks).

#### The PBMC10K dataset

is from the 10X Genomics and is annotated with 14 cell types. To align with the baseline model CFGen, we used the same preprocess procedure as CFGen, filtering out genes that are expressed in less than 20 cells. This remained 10,000 cells across 25,604 genes and 40,086 peaks.

#### The BABEL dataset

is a combination of four different datasets (DM, HSR, PBMC human data, and GM12878). Same as the BABEL, we removed genes encoded on sex chromosomes and cells expressing fewer than 200 genes or more than 7,000 genes. For peaks, we removed peaks on sex chromosomes, merged overlapping peaks, and removed peaks occurring in fewer than five cells or more than 10% of cells. This remained 40,465 paired data with 34,861 genes and 223,897 peaks.

#### The MiniAtlas dataset

is latest multi-omics atlas dataset with around 133,549 paired scRNA-seq data and scATAC-data. It has 36,601 genes and 524,151 peaks. We filtered the peaks that opened in less than 0.2% of all cells, remaining 165,564 peaks. The dataset includes cells from 19 tissues and 56 cell types, which makes it a suitable dataset for model pretraining.

For all the datasets, since scDiffusion-X can directly model count data, we did not perform data transformation (for example, logarithmic transformation) on the cell-by-feature count matrices. The scATAC data were binarized in all datasets. The Openproblem dataset and PBMC10k dataset were split randomly into 80% and 20% for training and validation. The BABEL dataset was split based on the Leiden cluster as stated in the previous section.

In the experiment of generating realistic single-cell multi-omics data, to make the scDesign3 available for comparison and evaluate the scalability of scDiffusion-X, we downsample the Openproblem dataset to form different size training data. Specifically, we selected the top 2,500 highly variable genes and peaks open in more than 30% of cells, which remained 5,542 peaks. We randomly select 5,000, 10,000, 20,000, and 40,000 cells to form training data with different sizes. In the experiment of perturbation translation, we selected the top 2,500 highly variable genes and peaks open in more than 15% of cells in the Openproblem dataset. To ensure the integrity of the information surrounding the target gene, we supplemented the dataset with peaks upstream and downstream of the target genes of 1M bp, resulting total 6,016 peaks.

### 4.9 Evaluation Metrics

Understanding the realism of generated cells and the biological information they contain is crucial. We developed metrics to evaluate the generated cells from both statistical and biological perspectives.

For the statistical aspect, we employed the Pearson Correlation Coefficient (PCC), Spearman Correlation Coefficient (SCC), Maximum Mean Discrepancy (MMD)^62^, and Local Inverse Simpson’s Index (LISI)^63^. The details of each metric are as follow:

#### Pearson Correlation Coefficient (PCC)

The PCC measures the strength of linear correlation between two vectors *x* and *y*. In our setting, *x* and *y* represent the average gene expression profiles of real and generated cells. It is defined as:

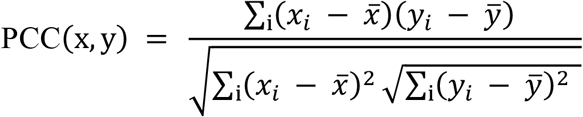

Where 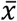and 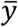are the means of *x* and *y*. The PCC ranges from -1 (perfect negative correlation) to 1 (perfect positive correlation), with 0 indicating no linear correlation.

#### Spearman Correlation Coefficient (SCC)

The SCC measures the strength of monotonic (rank-based) relationships between two variables (average gene expression profiles of real and generated cells in this case). Instead of raw values, it computes the PCC between the rank-transformed vectors:

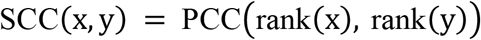

The SCC also ranges from -1 to 1, with higher values indicating stronger monotonic agreement.

#### Maximum Mean Discrepancy (MMD)

MMD quantifies the distance between two probability distributions (P) (real cells) and (Q) (generated cells) in a reproducing kernel Hilbert space (RKHS). Given kernel function *k*(⋅,⋅), the squared MMD is defined as:

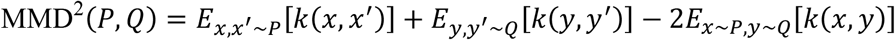

We computed MMD using the principal components of the real and generated cells. Smaller values indicate greater similarity between distributions, with 0 corresponding to identical distributions.

#### Local Inverse Simpson’s Index (LISI)

LISI quantifies how well real and generated cells are mixed in a shared neighborhood graph. Specifically, we first constructed a K-nearest neighbor (KNN) graph using 20 principal components and k=10 neighbors, and then applied the implementation from the scIB^64^ package (ilisi_graph). The score is based on the inverse Simpson’s index, which measures the diversity of real vs. generated labels in each cell’s neighborhood. The final LISI score is normalized to the range [0,1], where values closer to 1 indicate better local mixing (i.e., the generated cells are statistically indistinguishable from real cells), and values closer to 0 indicate poor mixing.

In terms of biological aspect, we included Marker Gene Kullback-Leibler Divergence (KLD), Random Forest Area Under the Curve (AUC), Uniform Manifold Approximation and Projection (UMAP) visualization^65^, and Pseudotime Correlation (PTC). The details of each metric are as follow:

#### Marker Gene Kullback–Leibler Divergence (KLD)

To assess whether the generated data preserve biologically meaningful signals, we focused on marker genes that define specific cell types. For each marker gene *g*, we computed its expression distribution in real cells (*P*_*g*_)and generated cells (*Q*_*g*_), and measured their divergence using Kullback–Leibler divergence:

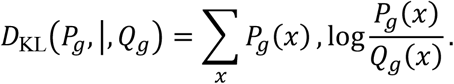

A smaller KLD value indicates that the generated cells closely match the real expression distribution of the marker gene, while larger values suggest distortion. We summarize results across all marker genes to evaluate fidelity.

#### Random Forest Area Under the Curve (AUC)

We trained a random forest classifier (1,000 trees, maximum depth = 5) to distinguish between real and generated cells, with data randomly split into 75% training and 25% validation sets. The discriminative performance was measured using the Receiver Operating Characteristic (ROC) Area Under the Curve (AUC):

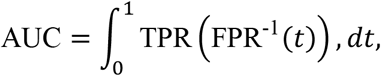

where TPR and FPR denote the true positive and false positive rates, respectively. An AUC close to 0.5 indicates that real and generated cells are indistinguishable, suggesting good biological realism, whereas an AUC closer to 1.0 indicates separability.

#### Uniform Manifold Approximation and Projection (UMAP)

We used UMAP to embed both real and generated cells into a shared two-dimensional space, enabling visual inspection of their global structure. While UMAP does not provide a numerical score, qualitative overlap of clusters and trajectories serves as evidence that generated data reproduce the biological heterogeneity of the real data.

#### Pseudotime Correlation (PTC)

To test whether developmental trajectories are preserved, we computed pseudotime values for each cell using Partition-based Graph Abstraction (PAGA).

We then measured the Pearson Correlation Coefficient (PCC) between the pseudotimes of real and generated cells:

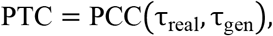

where τ denotes the pseudotime ordering. PTC values close to 1 indicate that the generated cells recapitulate the developmental progression observed in real data, while lower values suggest deviations.

In the evaluation of the DCA contribution, we used the information entropy (IE) and mean square error (MSE) to identify the most informative DCA and time step. For example, let 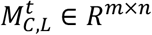represent the RNA to ATAC attention map for the C-th cell type at the t-th time step in the L-th DCA, where *n* and *m* are the RNA feature and ATAC feature dimensions. The IE is defined as:

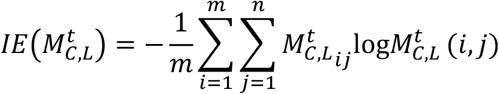

where 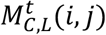stands for the i-th row and j-th column element of matrix 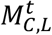. And the MSE is:

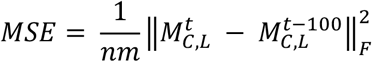

where ‖·‖_*F*_ denotes the Frobenius norm. Here the IE quantifies the information richness within a specific attention map, while the MSE assesses the change in information compared to the attention map from 100-time steps earlier.

### 4.10 Computational consumption

All experiments in our manuscript were conducted on a single NVIDIA 80G A100 GPU. We evaluated the computational resources and time for the model training and inference in Table S1. The small, normal, and large are three different model sizes corresponding to various data scales. We used the small setting for the scalability experiments with scDesign3, and the large setting for the modality translation. The remaining experiments are conducted under normal settings. We also evaluate the VAE-based model MultiVI as a comparison.

As shown in Table S1, the GPU consumption of scDiffusion-X is comparable to the classical bioinformatic model, which is small enough that most users can afford to use it. The training and inference time for the diffusion backbone is indeed longer than the simple VAE method. This is because denoising diffusion probability models have more complex model architectures and finer-grained generation processes, requiring more time to fit complex data distributions. As we mentioned in the Section 2.2, the additional computational cost is justified by the superior scalability and generation accuracy offered by diffusion-based methods. scDiffusion-X balances computational feasibility with modeling power, making it a practical and effective tool for large-scale single-cell multi-omics data generation and analysis.

## 5. Data availability

All data used in this paper are publicly available via the original publications and releases. The Openproblem dataset was downloaded from Gene Expression Omnibus (GEO), accession number GSE194122. The PBMC10k is available from the 10X company website: https://www.10xgenomics.com/support/single-cell-multiome-atac-plus-gene-expression. The BABEL dataset was downloaded from their official storage: https://drive.google.com/file/d/1J-4HH5e8rYapq5JtRq7G-foDs6Y9NzyQ/view. It contains the following datasets, which are all publicly available: Human data jointly profiling PBMC cells’ expression and chromatin accessibility is available from 10x Genomics’ data portal (https://support.10xgenomics.com/single-cell-multiome-atac-gex/datasets/1.0.0/pbmc_granulocyte_sorted_10k). Human data jointly profiling DM and HSR cells’ expression and chromatin accessibility is available through GEO (accession no. GSE160148). The external dataset Human data jointly profiling GM12878 cells is available through GEO (accession no. GSE166797). The ENCODE H3K27ac data is available at the ENCODE portal: https://www.encodeproject.org/. The chromatin loop data from the HiChIP database is available at the HiChIPdb: https://health.tsinghua.edu.cn/hichipdb/. The cross-modality perturbation data can be downloaded from PerturBase^67^: http://www.perturbase.cn/browser/19. The real multiome perturbation dataset is available at Zenodo:https://doi.org/10.5281/zenodo.15116138.

## 6. Code availability

The codes for implementing scDiffusion-X and for reproducing the results in this paper are available at the GitHub repository https://github.com/EperLuo/scDiffusion-X. The detailed instructions for scDiffusion-X are available at https://scDiffusionX.readthedocs.io/. The pretrained model weights and pretrained data are available at https://figshare.com/articles/dataset/scDiffusion-X/28582061.

## 7. Acknowledgements

The work is sponsored by The National Key R&D Program of China (2025YFC3409300), Tsinghua-Toyota Joint Research Institute Inter-disciplinary Program (20243930093), and National Natural Science Foundation of China (92470105, 62373210)

## Supplementary

**Figure S1.**
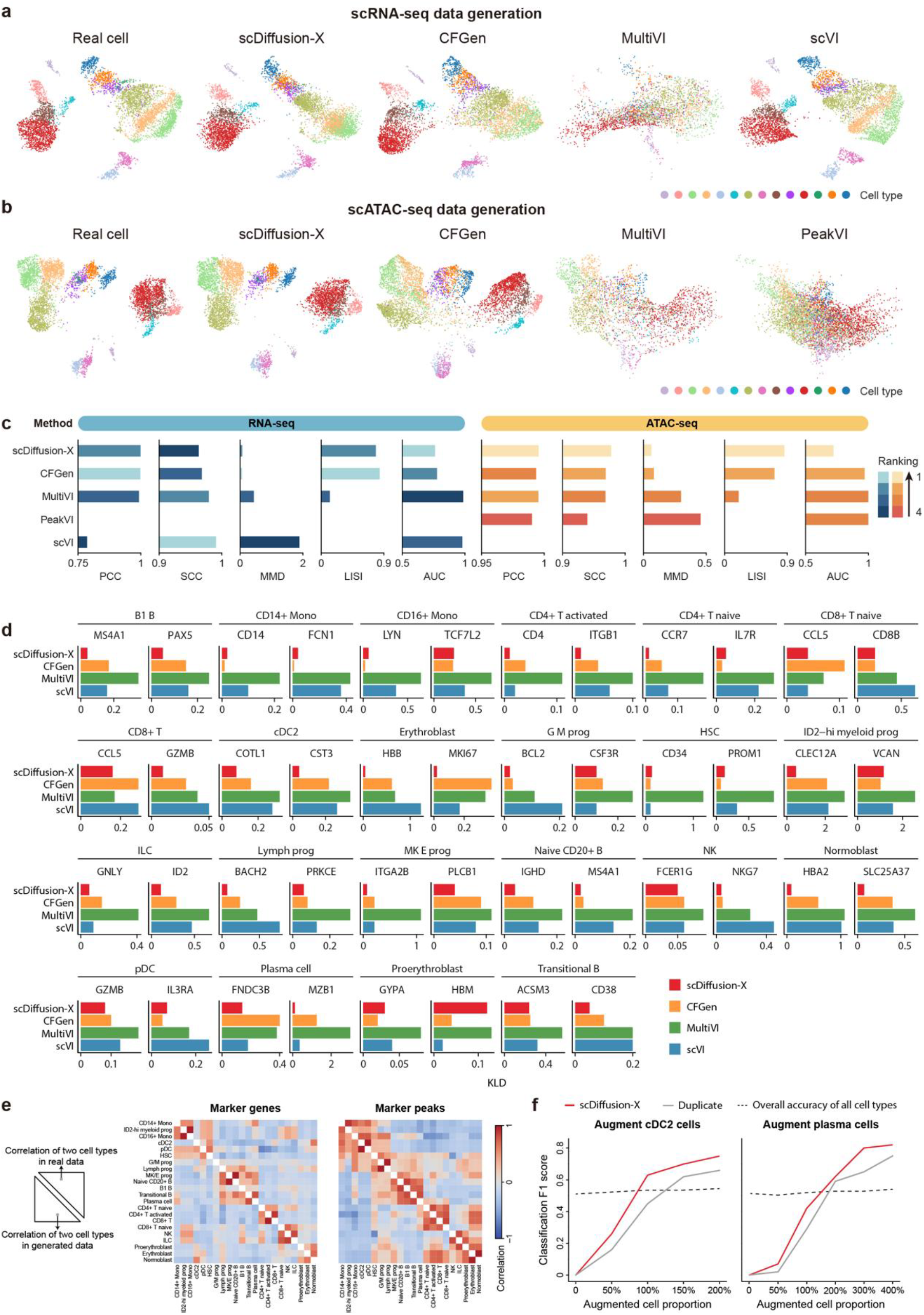
(a) The UMAP of generated scRNA-seq data by different methods on the PBMC10k dataset with pseudotime. (b) The UMAP of generated scATAC-seq data by different methods on the PBMC10k dataset. (c) Performance of different methods on the PBMC10k dataset under different evaluation metrics. (d) The KLD score of Marker gene expressions between real and generated cells in different cell types. For each cell type, we choose two marker genes. (e) The Pearson correlation score heatmap of the marker gene (and marker peak) within the real cell (upper right) and the heatmap within the generated cell (lower left). For each cell type, we picked 200 cells randomly and obtained the value of the top 1 differential expression gene (peak), which formed a *R*^1×200^ vector. We then calculated the Pearson correlation score between the vectors of each cell type. We did this in both real and generated cells to see if the generated cells can reproduce the relationship between different cell type markers. The generated cell’s heatmap (lower left) showed a very similar pattern to the real cell’s heatmap (upper right). (f) The effect of data augmentation on cell type classifiers. Here, Duplicate refers to duplicating the existing cell to balance its number on the original dataset. The classifier’s F1 scores for these two rare cell types increased significantly as the proportion of argument cells increased, showing more effectiveness than the data balancing method. The proportion here is the number of argument cells relative to the original number of cells.

**Figure S2.**
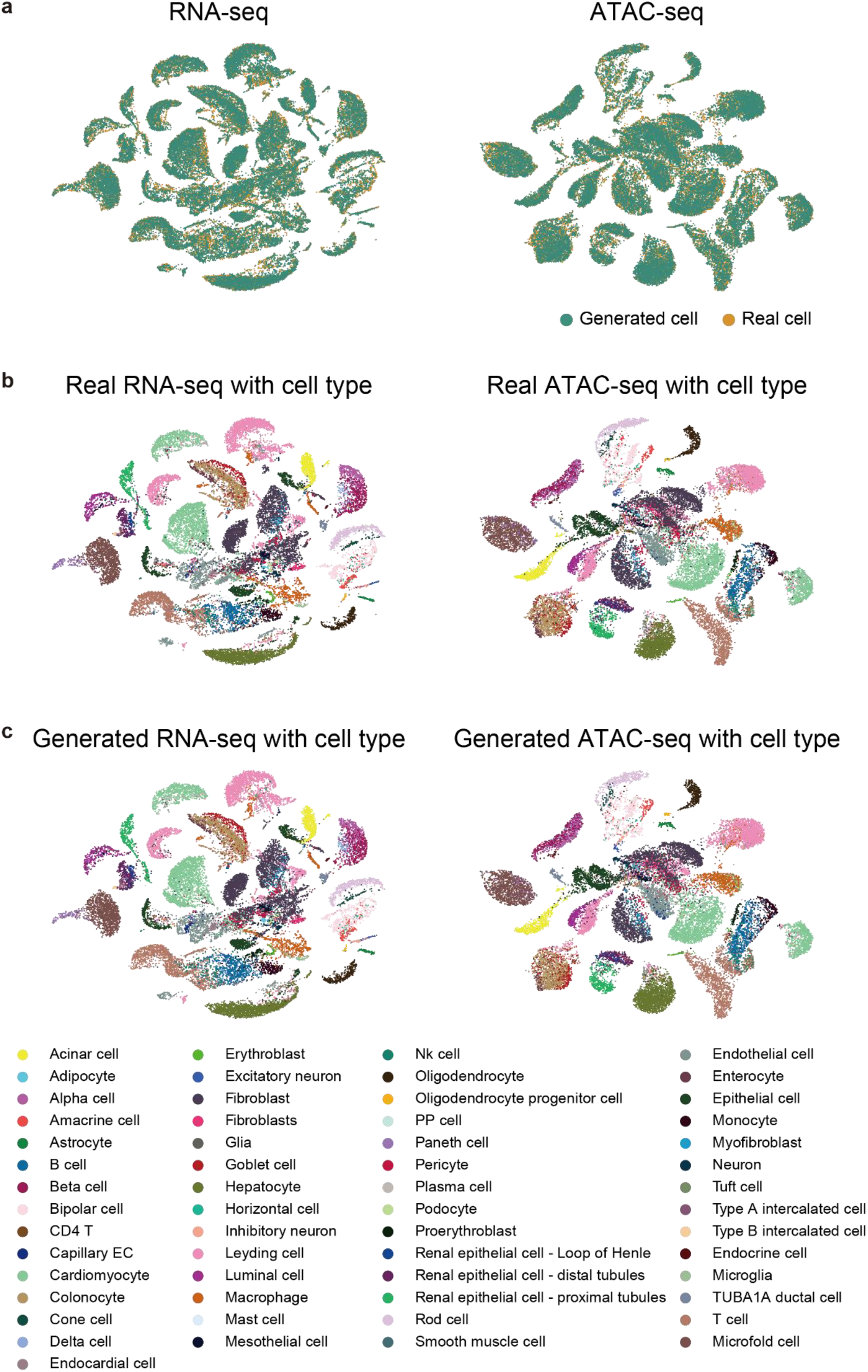
(a) The RNA-seq and ATAC-seq UMAP of real and generated cells in the MiniAtlas dataset. We conditionally generated each type of cells according to their cell numbers in the MiniAtlas dataset. (b) The RNA-seq and ATAC-seq UMAP of real cells colored with cell type in the MiniAtlas dataset. (c) The RNA-seq and ATAC-seq UMAP of scDiffusion-X conditionally generated cells colored with cell type in the MiniAtlas dataset.

**Figure S3.**
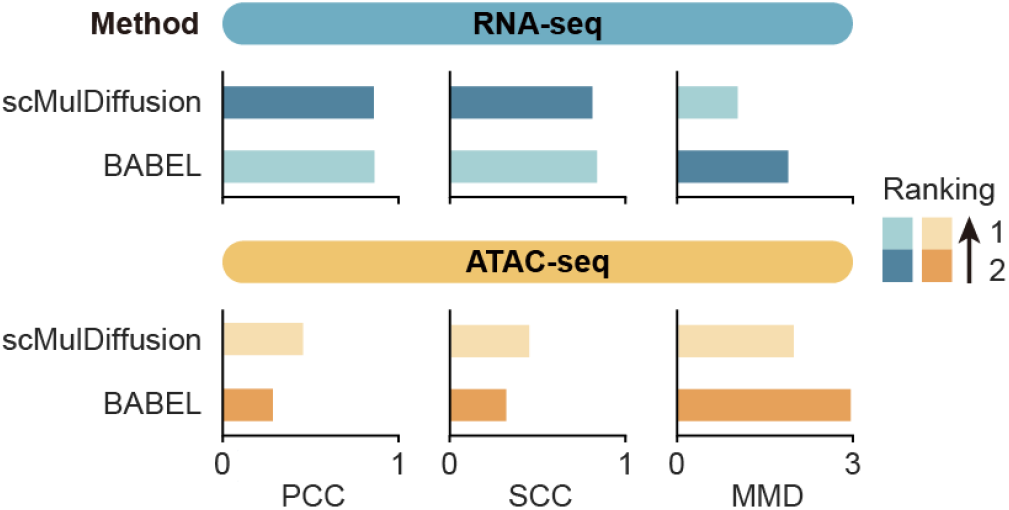
Performance of scDiffusion-X and BABEL translated external data (GM12878) under different evaluation metrics.

**Figure S4.**
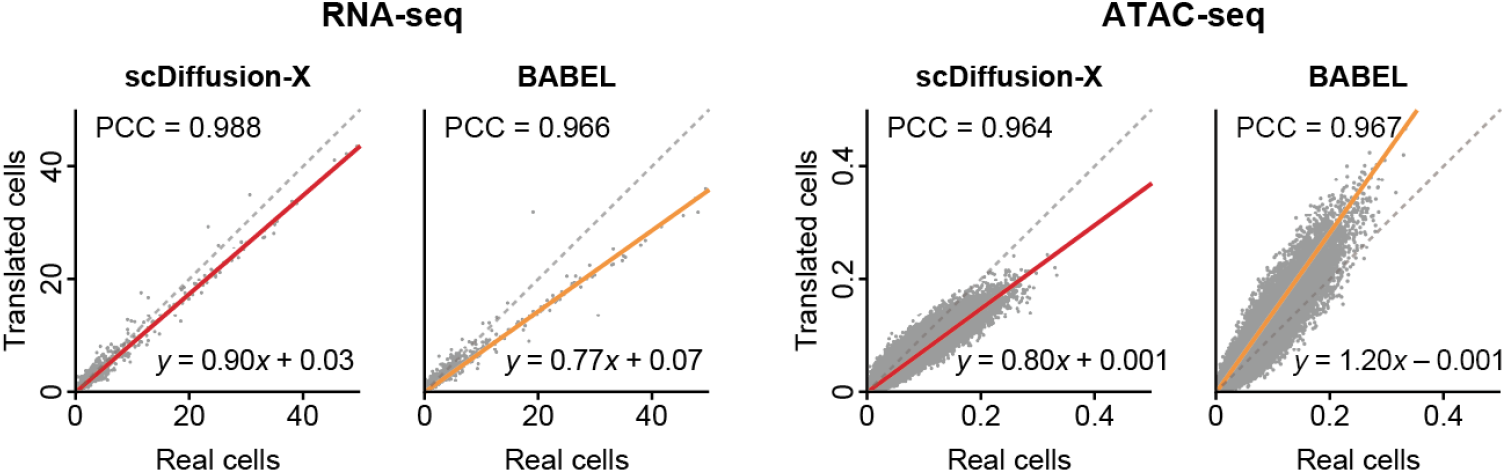
The comparison between pseudo bulk gene expression and chromatin accessibility using generated cells and real cells in the test set of BABEL dataset. Each dot represents a gene or peak.

**Figure S5.**
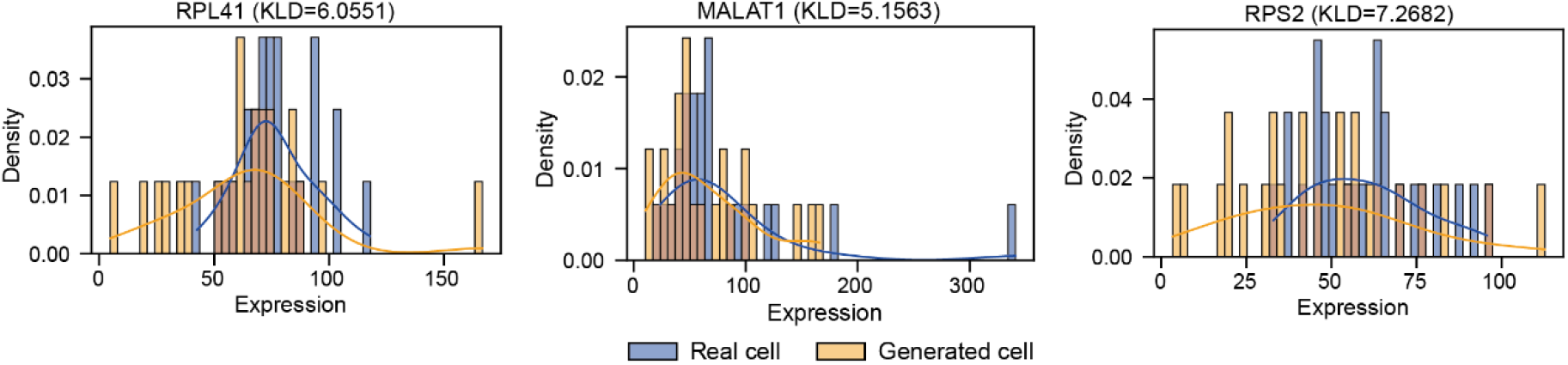
The gene expression distributions in the translation uncertainty experiments. These genes are the top 3 expressed genes in the real cell of the specific cell type. The distribution of these genes in real cells of this specific cell type and cells generated multiple times from a single cell shows a very similar pattern.

**Figure S6.**
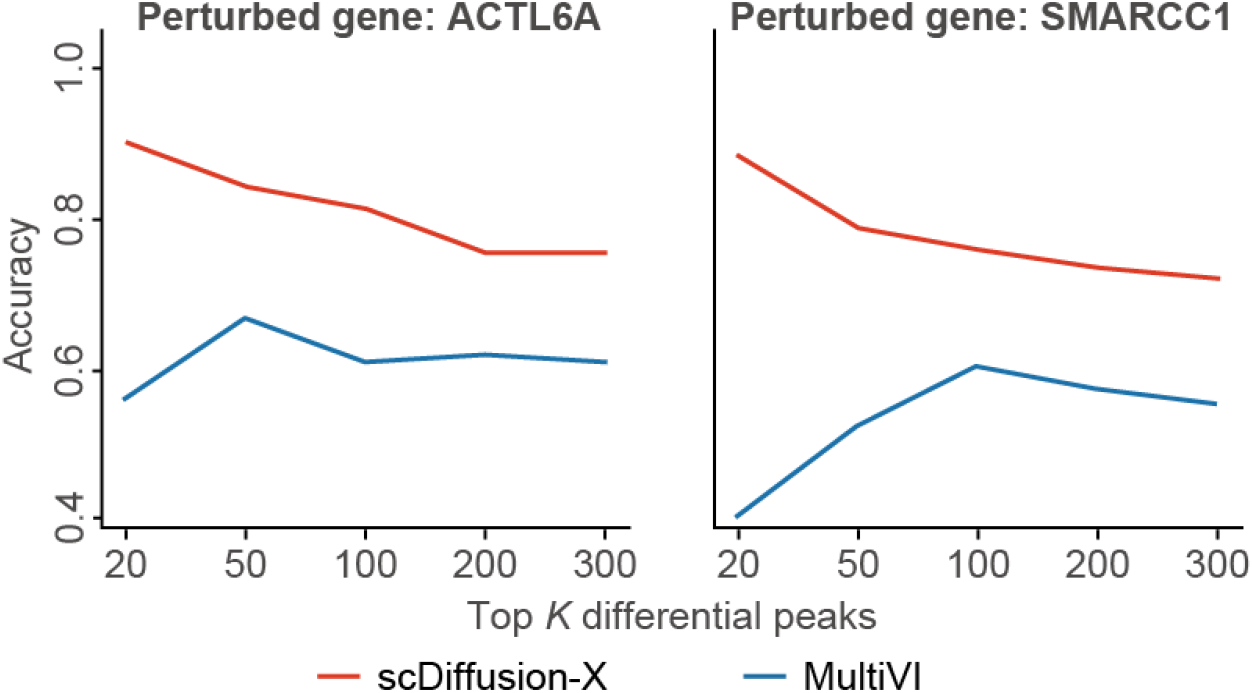
scDiffusion-X can reconstruct the key biological signal changes in real multiome perturbed scATAC-seq data. We used a real multiome perturbation dataset^46^ to test the model’s ability for cross-modality perturbation. The dataset applies Multiome Perturb-seq in a CRISPRi screen of 13 chromatin remodelers in human RPE-1 cells. We selected two perturbed genes (SMARCC1 and ACTL6A) as the test set and the rest of the perturbed genes and the control group as the training set to train scDiffusion-X. As a result, there are 4227 cells in the training set, including 1144 control cells and 3083 cells for 11 perturbations, and 497 cells in the test set for SMARCC1 and ACTL6A perturbations. We first merged the chromosome regions from 500-bp intervals (the original interval width in the dataset) into 1000-bp intervals. We then filtered 2500 highly variable genes and chromatin regions that open in more than 15% cells. We trained *scDiffusion-X* using cells in the training set, and translated perturbed scRNA-seq data in the test set into corresponding scATAC-seq data. Specifically, in the perturbation inference phase, we used the real perturbed scRNA-seq data as input and translated it into corresponding scATAC-seq data. We then calculated the change direction of predicted perturbed scATAC-seq data with the real control scATAC-seq data. We used the same setting to train and test the MultiVI model as a comparison. As shown in the figure Rx, scDiffusion-X achieved more than 80% accuracy in the top 100 differential peaks when perturbing the ACTL6A gene, and more than 70% accuracy in the top 100differential peaks when perturbing the SMARCC1 gene, both were higher than MultiVI.

**Figure S7.**
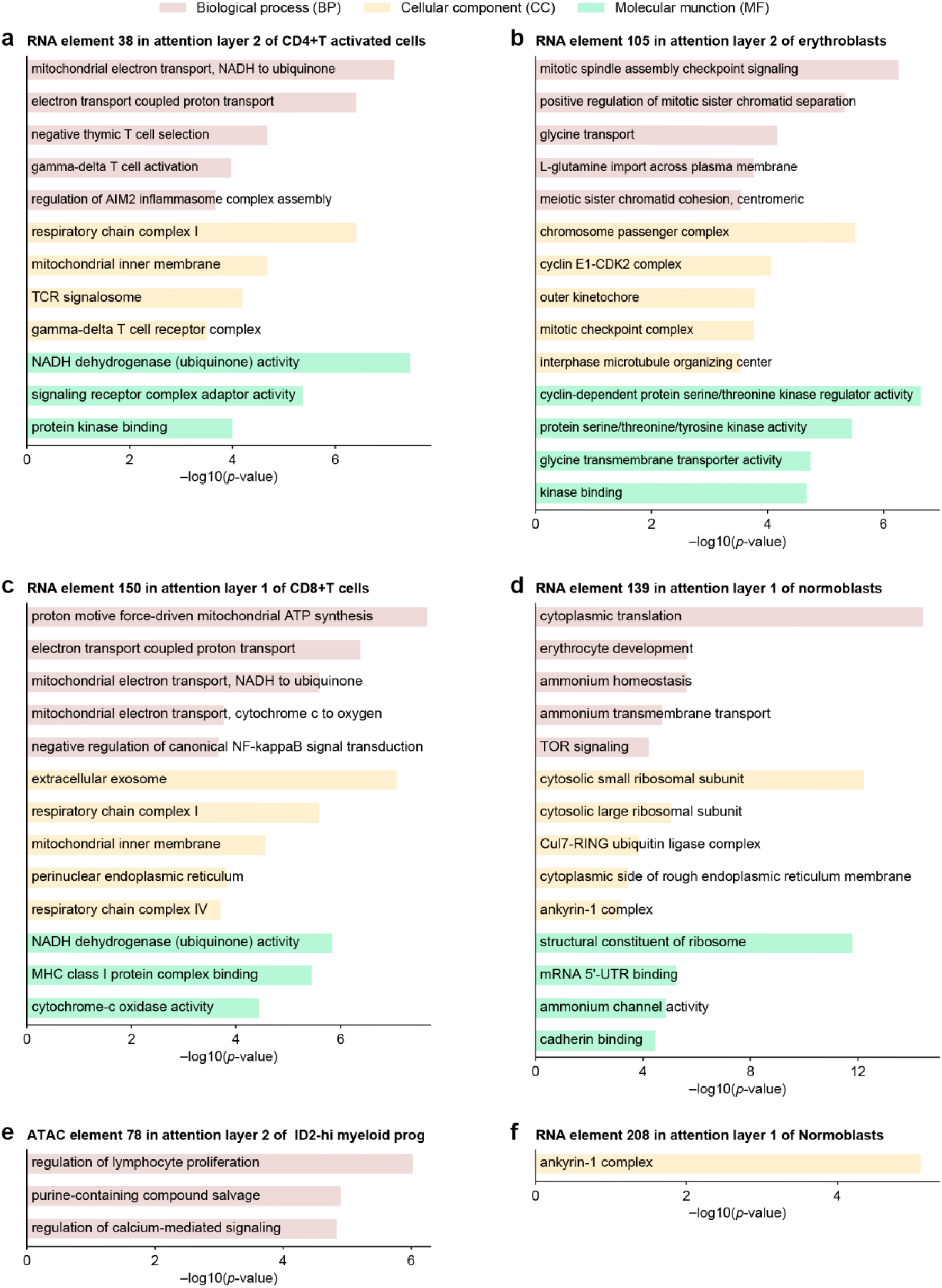
(a-b) GO terms of the genes associated with the 38th RNA elements in the second DCA for the CD4+T activated cell and GO terms of the genes associated with the 105th RNA elements in the second DCA for the Erythroblast cell. (c-d) GO terms of the genes associated with the 150th RNA elements in the first DCA for the CD8+T cell and GO terms of the genes associated with the 139th RNA elements in the first DCA for the Normoblast cell. (e-f) GO terms of the genes associated with the 78th ATAC elements in the second DCA for the ID2-hi myeloid prog cell and GO terms of the genes associated with the 208th ATAC elements in the first DCA for the Normoblast cell. Here we used the nearest gene of each key peak to obtain the GO terms. The GO terms related to these nearest genes are much fewer than those of key genes. We think this is because the key peak does not directly correspond to a specific gene, and the nearest genes of the key peaks set can not fully represent the function of these key peaks.

**Figure S8.**
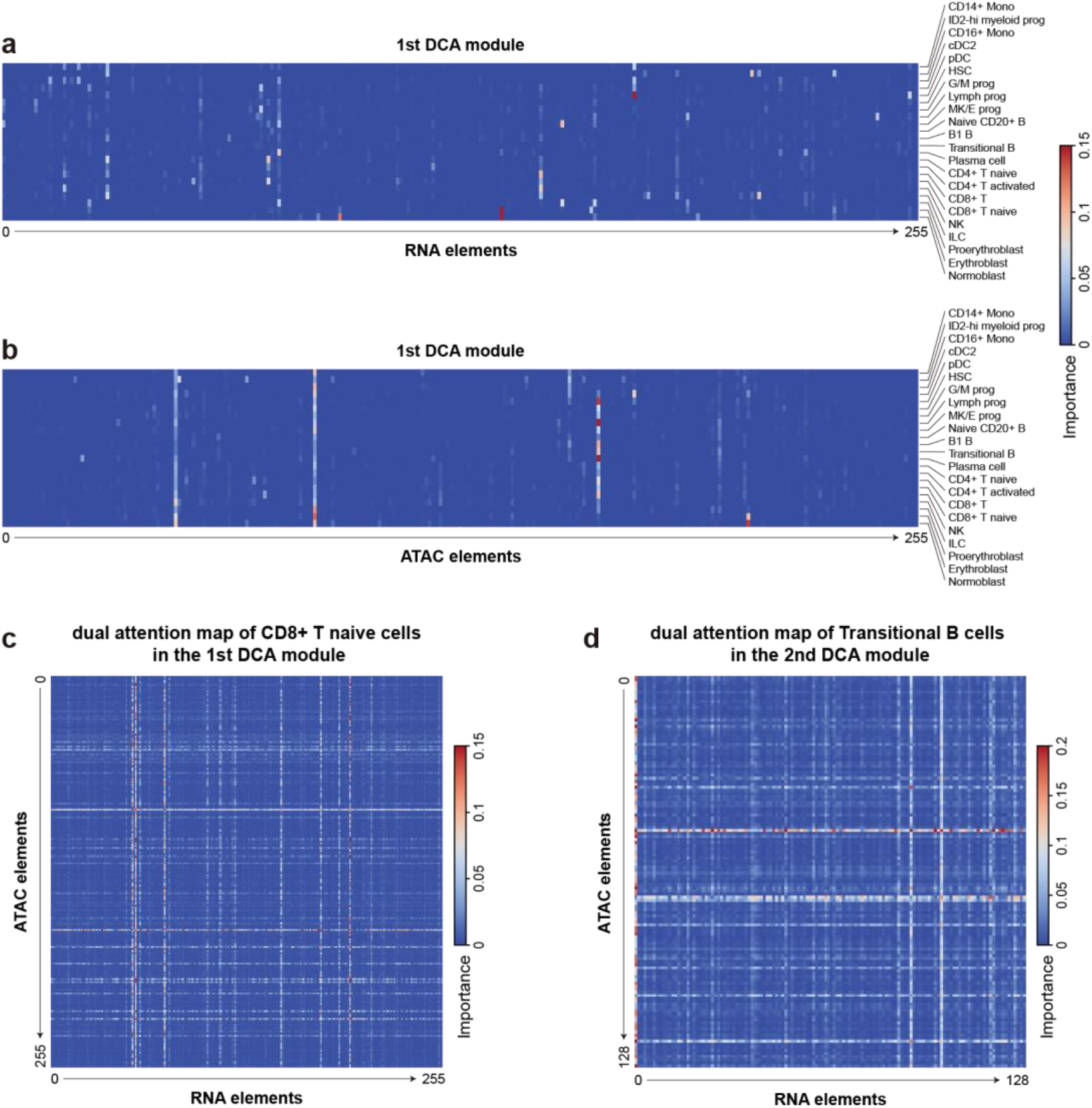
(a) Attention map of each RNA element to all ATAC elements in different cell types in the first DCA. (b) Attention map of each ATAC element to all RNA elements in different cell types in the first DCA. (c) Attention map of CD8+ T naive cell in the first DCA. (d) Attention map of Transitional B cell in the second DCA.

**Figure S9.**
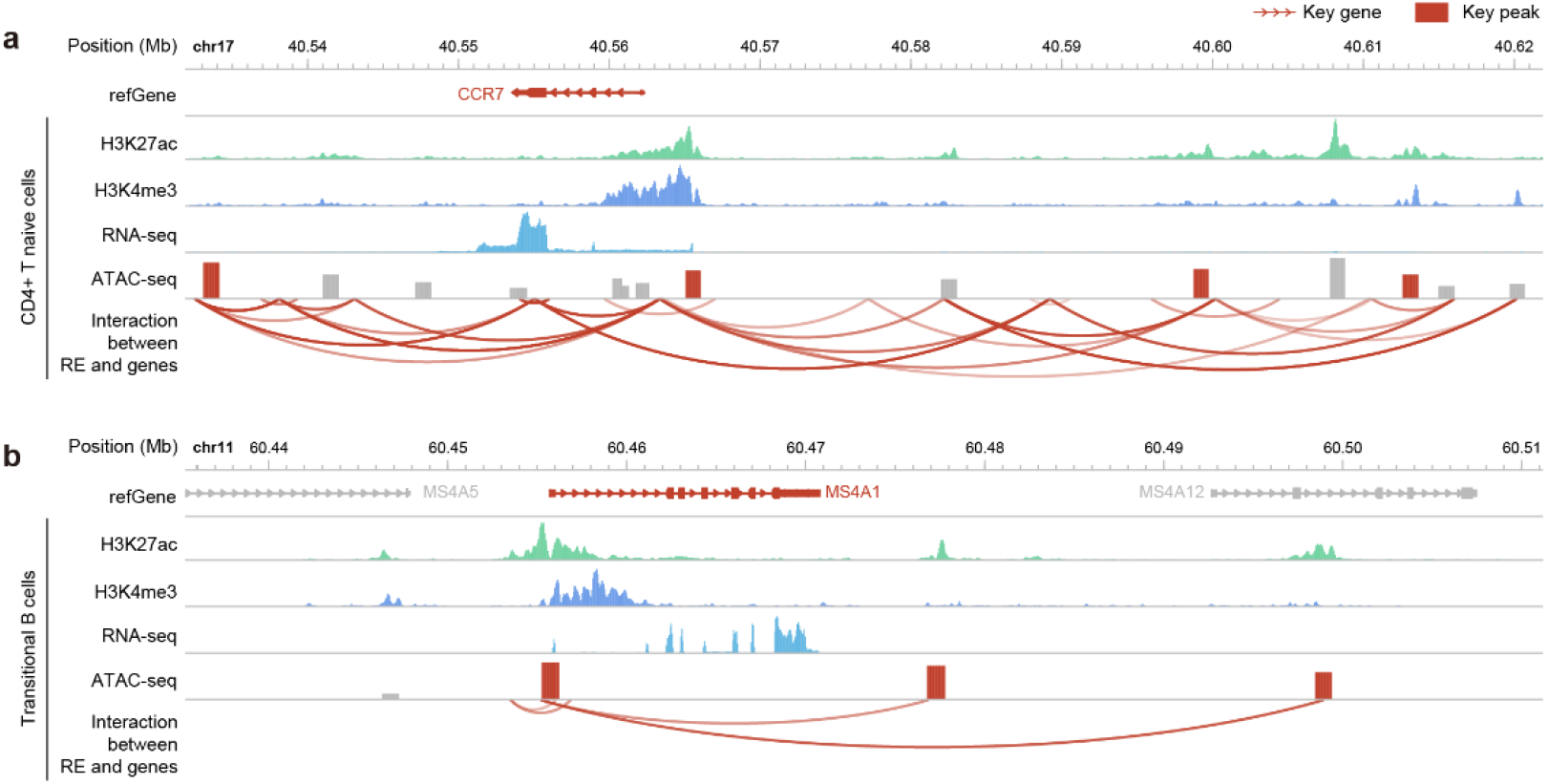
(a) A showcase of key genes and the key peaks in CD8+ T naive cell selected by scDiffusion-X. The key peaks are located in the potential enhancer position and tss position of the key gene CCR7. (b) A showcase of key genes and the key peaks in Transitional B cell selected by scDiffusion-X. The key peaks are located in the potential enhancer position and tss position of the key gene MS4A1.

**Figure S10.**
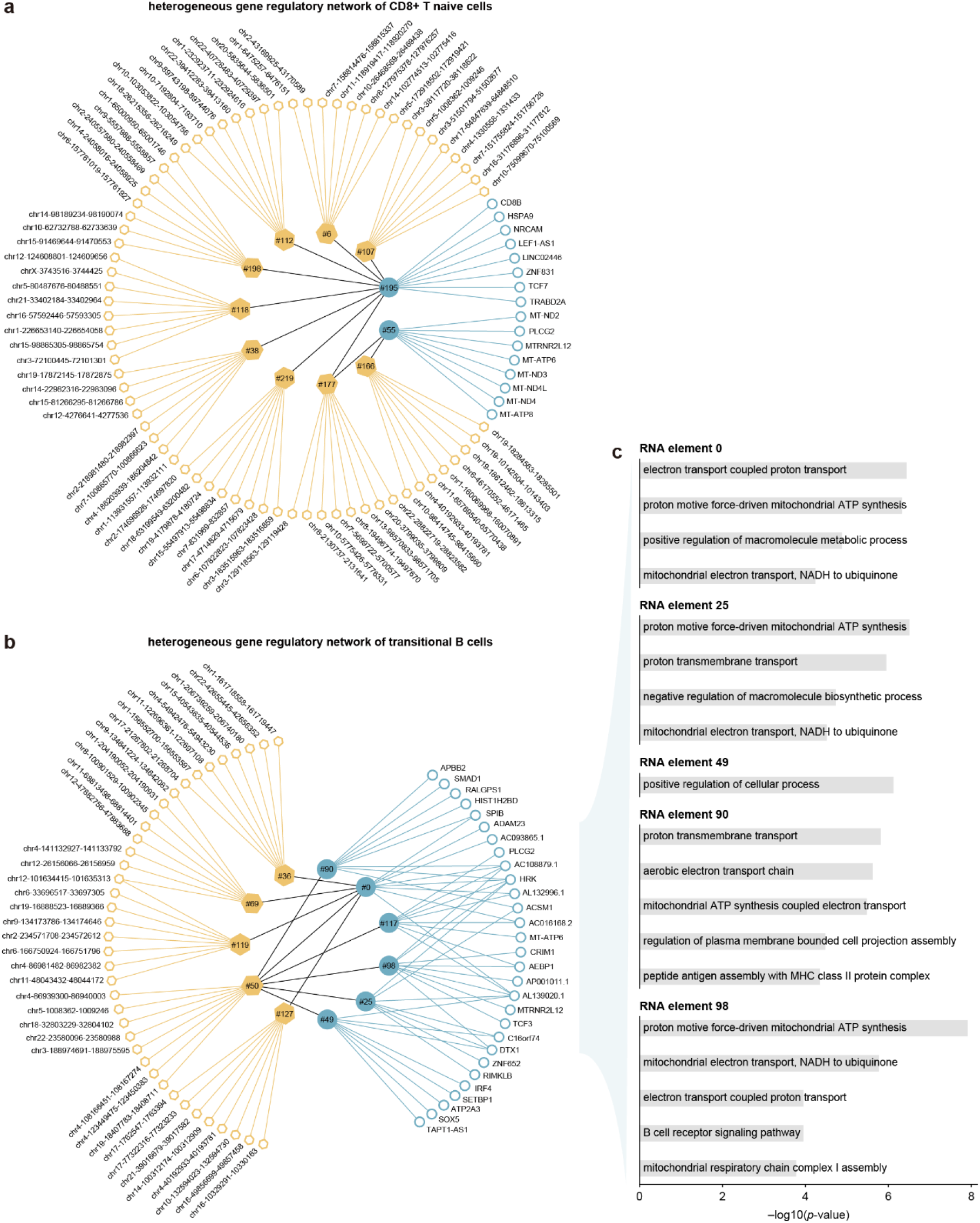
(a) A showcase of the CD8+ T naive cell heterogeneous network obtained by scDiffusion-X. This network shows the relationship of key genes – key RNA elements – key ATAC elements – key peaks. (b) A showcase of the Transitional B cell heterogeneous network obtained by scDiffusion-X. This network shows the relationship of key genes – key RNA elements – key ATAC elements – key peak. (c) The GO terms obtained from the key genes of each RNA element in the heterogeneous network.

**Figure S11.**
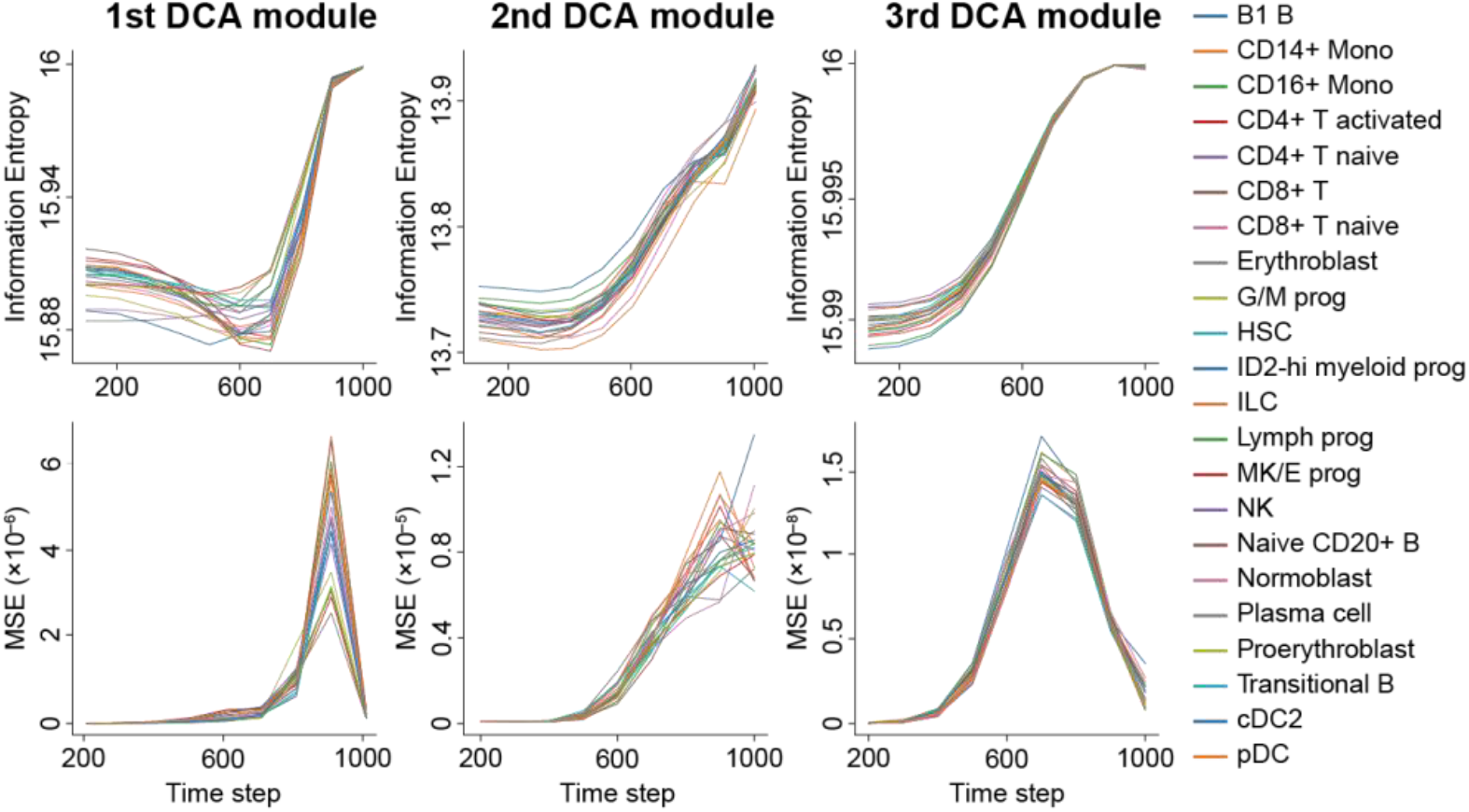
Investigating the effectiveness of the DCA. Ablation study for information richness of RNA to ATAC attention maps in different DCAs and different time steps.

**Figure S12.**
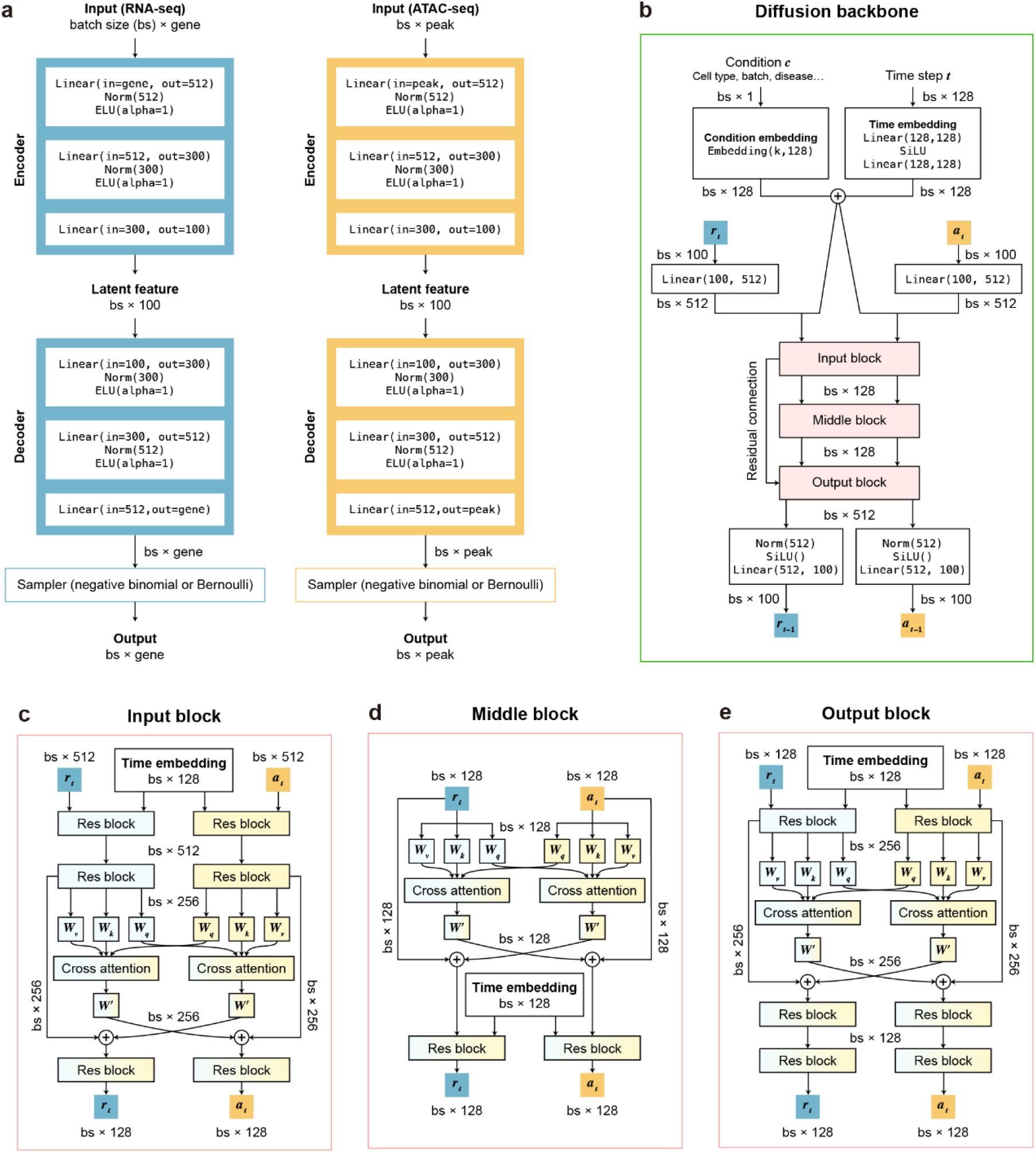
Model structure details. (a) Autoencoder structure and parameters details. (b) Multimodal denoising network structure and parameters details. (c) Input block structure and parameters details. (d) Middle block structure and parameters details. (e) Output block structure and parameters details.

**Table S1.** The computational resource and time required for different sizes of model.

| Model size | Model | Inference<br>time / 2000<br>cells | Training<br>time | GPU<br>consumption /<br>inference 2000<br>cell | GPU<br>consumption /<br>training batch<br>size 128 |
| --- | --- | --- | --- | --- | --- |
| Small | AE | 0.254 | 1.59 h | 948 MB | 1800 MB |
|  | Diffusion | 26.87 s | 6.66h | 4444 MB | 1498 MB |
| Normal | AE | 0.304 s | 1.84 h | 1474 MB | 2558 MB |
|  | Diffusion | 26.89 | 6.68 h | 4454 MB | 2024 MB |
| Large | AE | 0.391 s | 2.02 h | 2252 MB | 4514 MB |
|  | Diffusion | 26.95 | 6.71 h | 4480 MB | 3074 MB |
| Standard | MultiVI | 2.3 s | 1.6 h | 794 MB | 1504 MB |

**Table S2.** The precision (percentage of validated pairs) of different methods in finding potential regulatory relationships.

| Method | Number hit | precision |
| --- | --- | --- |
| ScDiffusion-X DCA | 237 | **0.6475** |
| Random pair | 119 | 0.3251 |
| Upregulated genes+ Nearest peak | 186 | 0.5167 |
| RE + nearest gene | 191 | 0.5219 |
| MultiVI+correlation | 189 | 0.5164 |
| SCENIC+ | 190 | 0.5191 |

## Notes

### Competing Interest Statement

The authors have declared no competing interest.

### Summary of Updates

Some formatting adjustments to the last submitted version

